# CORSID enables *de novo* identification of transcription regulatory sequences and genes in coronaviruses

**DOI:** 10.1101/2021.11.10.468129

**Authors:** Chuanyi Zhang, Palash Sashittal, Mohammed El-Kebir

## Abstract

Genes in coronaviruses are preceded by transcription regulatory sequences (TRSs), which play a critical role in gene expression mediated by the viral RNA-dependent RNA-polymerase via the process of discontinuous transcription. In addition to being crucial for our understanding of the regulation and expression of coronavirus genes, we demonstrate for the first time how TRSs can be leveraged to identify gene locations in the coronavirus genome. To that end, we formulate the TRS AND Gene Identification (TRS-Gene-ID) problem of simultaneously identifying TRS sites and gene locations in unannotated coronavirus genomes. We introduce CORSID (CORe Sequence IDentifier), a computational tool to solve this problem. We also present CORSID-A, which solves a constrained version of the TRS-Gene-ID problem, the TRS Identification (TRS-ID) problem, identifying TRS sites in a coronavirus genome with specified gene annotations. We show that CORSID-A outperforms existing motif-based methods in identifying TRS sites in coronaviruses and that CORSID outperforms state-of-the-art gene finding methods in finding genes in coronavirus genomes. We demonstrate that CORSID enables *de novo* identification of TRS sites and genes in previously unannotated coronaviruses. CORSID is the first method to perform accurate and simultaneous identification of TRS sites and genes in coronavirus genomes without the use of any prior information.

## 1 Introduction

Coronaviruses are comprised of a single-stranded RNA genome that is ready to be translated by the host ribosome (Fig. 1a). While the majority of messenger RNA (mRNA) in eukaryotes is *monocistronic, i.e*. each mRNA is translated into a single gene product, the coronavirus RNA genome is comprised of many genes, which are expressed and translated using two distinct mechanisms [15]. First, upon entry, the viral genome is translated to produce polypeptides corresponding to one or two overlapping open reading frames (ORFs). Second, the resulting polypeptides undergo auto-cleavage, producing many non-structural proteins, including RNA-dependent-RNA-polymerase (RdRP), which mediates the expression of the remaining viral genes via *discontinuous transcription* [22]. That is, RdRP is prone to perform template switching upon encountering *transcription regulatory sequences* (TRSs) located in the 5’ untranslated region (UTR) of the genome — called TRS-L where L stands for leader — and upstream of each viral gene — called TRS-B where B stands for body (Fig. 1b). This mechanism yields multiple subgenomic mRNAs that are translated into the structural and accessory viral proteins, necessary for the viral life cycle. Not only is the identification and characterization of TRS sites crucial to understanding the regulation and expression of the viral proteins, but here we hypothesize that the existence of these regulatory sequences can be leveraged to simultaneously identify TRS sites and associated viral genes in unannotated coronavirus genomes with high accuracy.

**Figure 1:**
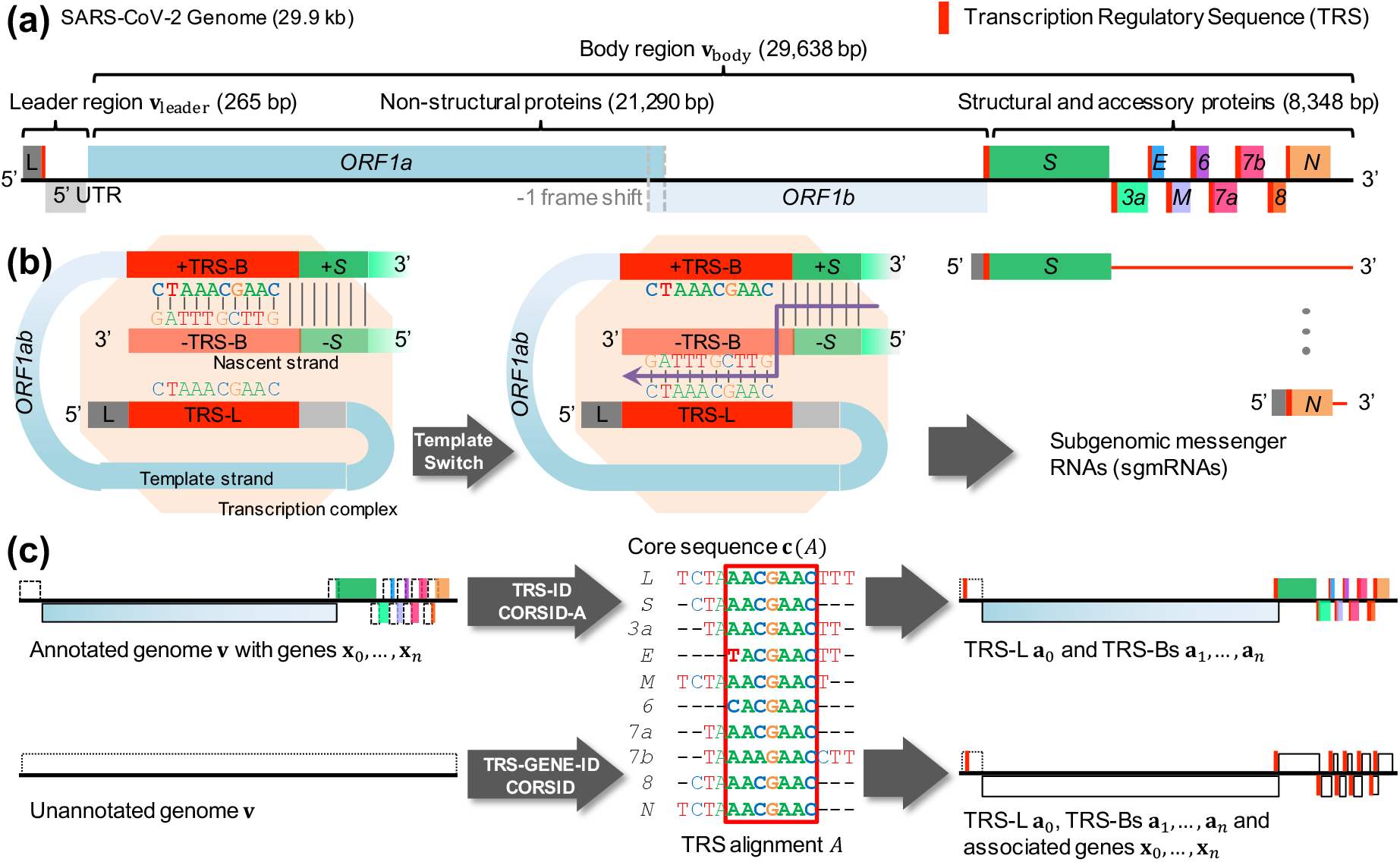
Overview. (a) A coronavirus genome **v** consists of a leader region **v**_leader_ and a body region **v**_body_. (b) Structural and accessory genes are expressed via discontinuous transcription with template switching occurring at transcription regulatory sequences (TRS, indicated in red), resulting in subgenomic messenger RNAs (sgmRNAs) for each gene. (c) In the TRS Identification (TRS-ID) problem, we wish to identify TRSs given a genome **v** with genes **x**_0_,*…*, **x**_*n*_. The TRS and Gene Identification (TRS-Gene-ID) asks to simultaneously identify genes and their associated TRSs given genome **v**. Throughout this manuscript we use ‘T’ (thymine) rather than ‘U’ (uracil).

While there exist methods for identifying either TRS sites or viral genes, no method exists that does so simultaneously (Table S1). More specifically, since TRSs contain 6 − 7 nt long conserved sequences, called *core sequences* [8,25], general-purpose motif finding methods [2,7,20,29] can be employed to identify TRS-L and TRS-Bs in coronaviruses. For instance, MEME [2] is a widely used method that employs expectation maximization to identify multiple appearances of multiple motifs simultaneously. The only method that is specifically developed for identifying TRS sites in coronaviruses is SuPER [28], which takes as input a coronavirus genome sequence with specified gene locations as well as additional taxonomic and secondary structure information. Importantly, SuPER as well as other general-purpose motif finding algorithms are unable to identify viral genes in unannotated coronavirus genome sequences. On the other hand, gene prediction is a well-studied problem with many methods including Glimmer3 [5, 21] and Prodigal [12, 13]. Glimmer3 uses a Markov model to assign scores to ORFs, and then processes overlapping genes to generate the final list of predicted genes. By contrast, Prodigal employs a more heuristic approach with fine-tuned parameters that are optimized to identify genes in prokaryotes. However, these general-purpose gene finding tools are not designed to leverage the genomic structure of coronaviruses, specifically the TRS sites located upstream of the genes in the genome, nor are they able to directly identify these regulatory sequences.

In this study, we introduce the TRS Identification (TRS-ID) and the TRS and Gene Identi-fication (TRS-Gene-ID) problems, to identify TRS sites in a coronavirus genome with specified gene annotations, and to simultaneously identifying TRS sites and genes in an unannotated coronavirus genome, respectively (Fig. 1c). Underpinning our approach is the concept of a *TRS alignment*, which is a multiple sequence alignment of TRS sites with additional constraints that result from template switching by RdRP. We introduce CORSID-A, a dynamic programming (DP) algorithm to solve the TRS-ID problem, adapting the recurrence that underlies the Smith-Waterman algorithm [24] for local sequence alignment. Additionally, we introduce CORSID to solve the TRS-Gene-ID problem via a maximum-weight independent set problem [11] on an interval graph defined by the candidate ORFs in the genome with weights obtained from the previous DP. We evaluate the performance of our methods on 468 coronavirus genomes downloaded from GenBank, demonstrating that CORSID-A outperforms MEME and SuPER in identifying TRS sites and, unlike these methods, possesses the ability to identify recombination events. Moreover, we find that CORSID vastly outperforms state-of-the-art gene finding methods. Finally, we illustrate how CORSID enables *de novo* identification of TRS sites and genes in previously unannotated coronaviruses. In summary, CORSID is the first method to perform accurate and simultaneous identification of TRS sites and genes in coronavirus genomes without the use of prior taxonomic or secondary structure information.

## 2 Problem Statement

We begin by introducing notation and key definitions (Section 2.1), followed by stating the TRS Identifi-cation problem (Section 2.2) and then the TRS and Gene Identification problem (Section 2.3).

### 2.1 Preliminaries

A *genome* **v** = *v*_1_ *… v*_|**v**|_ is a sequence from the alphabet Σ = {A, T, C, G}. The first position of the genome is known as the *5’ end* whereas the last position of the genome is known as the *3’ end*. We denote the contiguous subsequence *v*_*p*_ *… v*_*q*_ of **v** by **v**[*p, q*]. We call a contiguous subsequence **x** of **v** also a *region*, denoted as **x** = [*x*^−^, *x*^+^] such that **x** = **v**[*x*^−^, *x*^+^]. Thus, coordinates *x*^−^ and *x*^+^ of a subsequence **x** are in terms of the reference genome **v**, *i.e*. 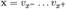 .Alternatively, we may refer to individual characters in a subsequence **x** using relative indices, *i.e*. **x** = *x*_1_ *… x*_|**x**|_. Our goals are twofold: given a coronavirus genome **v**, we aim to identify (i) TRS-L and TRS-Bs, and optionally, (ii) the associated genes (Fig. 1c). To begin, recall the following definition of an alignment.

#### Definition 1.

Matrix *A* = [*a*_*ij*_] with *n* + 1 rows is an *alignment* of sequences **w**_0_, *…*, **w**_*n*_ ∈ Σ^*^ provided (i) entries *a*_*ij*_ either correspond to a letter in the alphabet Σ or a gap denoted by ‘−’ such that (ii) no column of *A* is composed of only gaps, and (iii) the removal of gaps of row *i* of *A* yields sequence **w**_*i*_.

Here, we seek an alignment with two additional constraints, called a TRS alignment defined as follows.

#### Definition 2.

An alignment *A* = [**a**_0_, *…*, **a**_*n*_]^*T*^ is a *TRS alignment* provided (i) **a**_0_ does not contain any gaps, and (ii) **a**_1_, *…*, **a**_*n*_ do not contain any internal gaps.

Intuitively, the first sequence **a**_0_ in the alignment *A* represents TRS-L, whereas **a**_1_, *…*, **a**_*n*_ represent TRS-Bs, each upstream of an accessory or structural gene. We do not allow gaps in the TRS-L sequence **a**_0_ as template switching by RdRP occurs due to complementary base pairing between TRS-L and the nascent strand of TRS-B [26]. For the same reason, we do not allow internal gaps in TRS-Bs **a**_*i*_. However, as each TRS-B may match a different region of the TRS-L, we do allow flanking gaps in these sequences (Fig. 1c). We score a TRS alignment *A* using a scoring function *δ* : Σ × (Σ ∪ {−}) → ℝ in the following way.

#### Definition 3.

The *score s*(*A*) of a TRS alignment *A* = [**a**_0_, *…*, **a**_*n*_]^*T*^ is given by 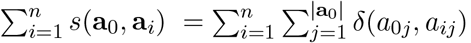, whereas the *minimum score s*_min_(*A*) is defined as min_*i*∈{1,…,*n*}_ *s*(**a**_0_, **a**_*i*_).

In other words, we score each TRS-B **a**_*i*_ (where *i* ≥ 1) by comparing it to the TRS-L sequence **a**_0_ in a way that is consistent with the mechanism of template switching during discontinuous transcription. As such, our scoring function differs from the traditional sum-of-pairs scoring function [3] where every unordered pair (**a**_*i*_, **a**_*j*_) of sequences contributes to the score of the alignment. Furthermore, each TRS alignment uniquely determines the core sequence as follows.

#### Definition 4.

Sequence **c**(*A*) is the *core sequence* of a TRS alignment *A* = [**a**_0_, *…*, **a**_*n*_]^*T*^ provided **c**(*A*) is the largest contiguous subsequence of **a**_0_ such that no character of **c** is aligned to a gap in any of **a**_1_, *…*, **a**_*n*_.

Note that the core sequence is a subsequence of the TRS sequences. As such, the TRS alignment can include nucleotides immediately flanking the core sequence, which have been shown to play an important role in discontinuous transcription in previous experiments [26].

### 2.2 The TRS IDENTIFICATION problem

The first problem we consider is that of identifying TRS sites given a viral genome with known genes **x**_0_, *…*, **x**_*n*_. Specifically, we are given a candidate region **w**_0_ that contains the unknown TRS-L **a**_0_ upstream of gene **x**_0_ as well as candidate regions **w**_1_, *…*, **w**_*n*_ that contain the unknown TRS-Bs **a**_1_, *…*, **a**_*n*_ of genes **x**_1_, *…*, **x**_*n*_. Section 3.1 details how to obtain these candidate regions when only given the gene locations. To further guide the optimization problem, we impose an additional constraint on the sought TRS alignment *A* in the form of a minimum length *ω* on the core sequence **c**(*A*) as well as a threshold *τ* on the minimum score *s*_min_(*A*) of the TRS alignment. We formalize this problem as follows.

**Problem 1** (TRS Identification (TRS-ID)). Given non-overlapping sequences **w**_0_, *…*, **w**_*n*_, core-sequence length *ω >* 0 and score threshold *τ >* 0, find a TRS alignment *A* = [**a**_0_, *…*, **a**_*n*_]^*T*^ such that (i) **a**_*i*_ corresponds to a subsequence in **w**_*i*_ for all *i* ∈ {0, *…, n*}, (ii) the core sequence **c**(*A*) has length at least *ω*, (iii) the minimum score *s*_min_(*A*) is at least *τ*, and (iv) the alignment has maximum score *s*(*A*).

### 2.3 The TRS AND GENE IDENTIFICATION problem

In the second problem, we are no longer given an annotated genome with gene locations. Rather, we seek to simultaneously identify genes and TRS sites given a viral genome sequence **v** split into a leader region **v**_leader_ and body region **v**_body_. Section 3.2 describes a heuristic for identifying these two regions when only given **v**. The key idea here is that each TRS alignment will uniquely determine a set of genes it encodes. To make this relationship clear, we begin by defining an open reading frame as follows.

#### Definition 5.

A contiguous subsequence **x** = [*x*^−^, *x*^+^] of **v** is an *open reading frame* provided **x** (i) has a length |**x**| that is a multiple of 3, (ii) starts with a start codon, *i.e. x*_1_ *… x*_3_ = ATG, and (iii) ends at a stop codon, *i.e. x*_|**x**|−2_ *… x*_|**x**|_ ∈ {ATG, TAG, TGA}.

Each TRS-B **a**_*i*_ is associated with at most one ORF that occurs immediately downstream of **a**_*i*_. Naively, to identify the ORF associated with **a**_*i*_, one could simply scan downstream of the TRS-B for the first occurrence of a start codon and continue scanning to identify the corresponding in-frame stop codon. However, this would not take ribosomal leaky scanning into account, where the ribosome does not initiate translation at the first encountered ‘ATG’. Section 3.2 provides a more robust definition of a downstream ORF that takes ribosomal leaky scanning into account. To summarize, we have that a TRS alignment *A* = [**a**_0_, *…*, **a**_*n*_]^*T*^ uniquely determines a set Γ(*A*) of candidate genes.

#### Definition 6.

A set Γ(*A*) of ORFs are *induced genes* of a TRS alignment *A* = [**a**_0_, *…*, **a**_*n*_]^*T*^ provided Γ(*A*) is composed of the ORFs that occur downstream of each TRS-B **a**_1_, *…*, **a**_*n*_ in **v**_body_.

Note that multiple TRS-Bs of a TRS alignment *A* = [**a**_0_, *…*, **a**_*n*_]^*T*^ can induce the same gene in **v**_body_. Moreover, there may *not* be an ORF downstream of a TRS-B **a**_*i*_. As such, we have that |Γ(*A*)| ≤ *n*. By contrast, in coronaviruses, each viral gene typically has a unique TRS-B. Moreover, these viral genes are typically non-overlapping in the genome. Finally, coronavirus genomes tend to be compact with most positions coding for genes. To capture these biological constraints, we introduce the following definitions.

#### Definition 7.

A TRS alignment *A* = [**a**_0_, *…*, **a**_*n*_]^*T*^ is *concordant* provided (i) each TRS-B **a**_*i*_ corresponds to exactly one gene in Γ(*A*), and (ii) there are no two ORFs in Γ(*A*) whose positions in **v**_body_ overlap.

#### Definition 8.

The *genome coverage g*(*A*) of a TRS alignment *A* is the number of positions in **v**_body_ that are covered by the set Γ(*A*) of induced genes.

This leads to the following problem.

**Problem 2** (TRS and Gene Identification (TRS-Gene-ID)). Given leader region **v**_leader_, body region **v**_body_, core-sequence length *ω >* 0 and score threshold *τ >* 0, find a TRS alignment *A* = [**a**_*i*_] such that (i) **a**_0_ corresponds to a subsequence in **v**_leader_, (ii) **a**_*i*_ corresponds to a subsequence in **v**_body_ for all *i* ≥ 1, (iii) the core sequence **c**(*A*) has length at least *ω*, (iv) the minimum score *s*_min_(*A*) is at least *τ*, (v) *A* is concordant, and (vi) *A* induces a set Γ(*A*) of genes with maximum genome coverage *g*(*A*) and *A* subsequently has maximum score *s*(*A*).

## 3 Methods

Section 3.1 introduces CORSID-A, which solves the TRS-ID problem. Section 3.2 introduces CORSID, solving the TRS-Gene-ID problem. Both sections discuss practical considerations as well as several heuristics for obtaining the required input to each problem. We implemented both methods in Python. The source code is available at https://github.com/elkebir-group/CORSID.

### 3.1 Solving the TRS IDENTIFICATION problem

Recall that in the TRS-ID problem we seek a TRS alignment *A* given input candidate regions sequences **w**_0_, *…*, **w**_*n*_ that each occur upstream of genes **x**_0_, *…*, **x**_*n*_. Intuitively, we define the candidate region for a gene **x**_*i*_ as the region 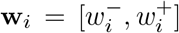 composed of positions *w*^−^ ≤ *p* ≤ *w*^+^ such that any sgmRNA starting at *p* will lead to the translation of ORF **x**_*i*_ by the ribosome. SuPER [28], the only other method for identifying TRSs in annotated coronavirus genomes, employs a heuristic by defining the candidate region **w**_*i*_ of a gene 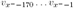, *i.e*. the candidate region **w**_*i*_ is a subsequence of 170 nt immediately upstream of gene **x**_*i*_ for 1 ≤ *i* ≤ *n*. Here, we take a more rigorous and flexible approach that takes ribosomal leaky scanning into account by skipping over previous ORFs with length smaller than 100 nt (details in Appendix A.1).

Recall that in a TRS alignment *A* = [**a**_0_, *…*, **a**_*n*_]^*T*^ only the TRS-Bs **a**_1_, *…*, **a**_*n*_ are allowed to have gaps (restricted to the flanks), and that the TRS-L **a**_0_ is gapless. To score a TRS alignment, we use a simple scoring function *δ* : Σ × (Σ ∪ {−}) → ℝ such that *s*(*x, y*) equals +1 for matches (*i.e. x* = *y*), − 2 for mismatches (*i.e. x ≠ y* and *y ≠* −), and 0 for gaps (*i.e. y* = −). In other words, while we reward matches and penalize mismatches, we do not penalize flanking gaps.

Recall that the sought TRS alignment *A* must induce a core sequence **c**(*A*) of length at least *ω*. Due to this constraint, the input sequences **w**_0_, *…*, **w**_*n*_ depend on one another and cannot be considered in isolation. We break this dependency by considering a subsequence **u** within **w**_0_ of length *ω*, restricting the induced core sequence **c**(*A*) of output TRS alignments *A* to contain **u**. We solve this constrained version of the TRS-ID problem using dynamic programming in time *O*(|**w**_0_|*L*) where *L* is the total length of candidate regions **w**_1_, *…*, **w**_*n*_ (details are in Appendix A.2 and Fig. 2a). We obtain the solution to the original TRS-ID problem by identifying the window **u** that induces a TRS alignment *A* with maximum score. As there are *O*(|**w**_0_|) windows in **w**_0_ of fixed length *ω*, this procedure takes *O*(|**w**_0_|^2^*L*) time.

**Figure 2:**
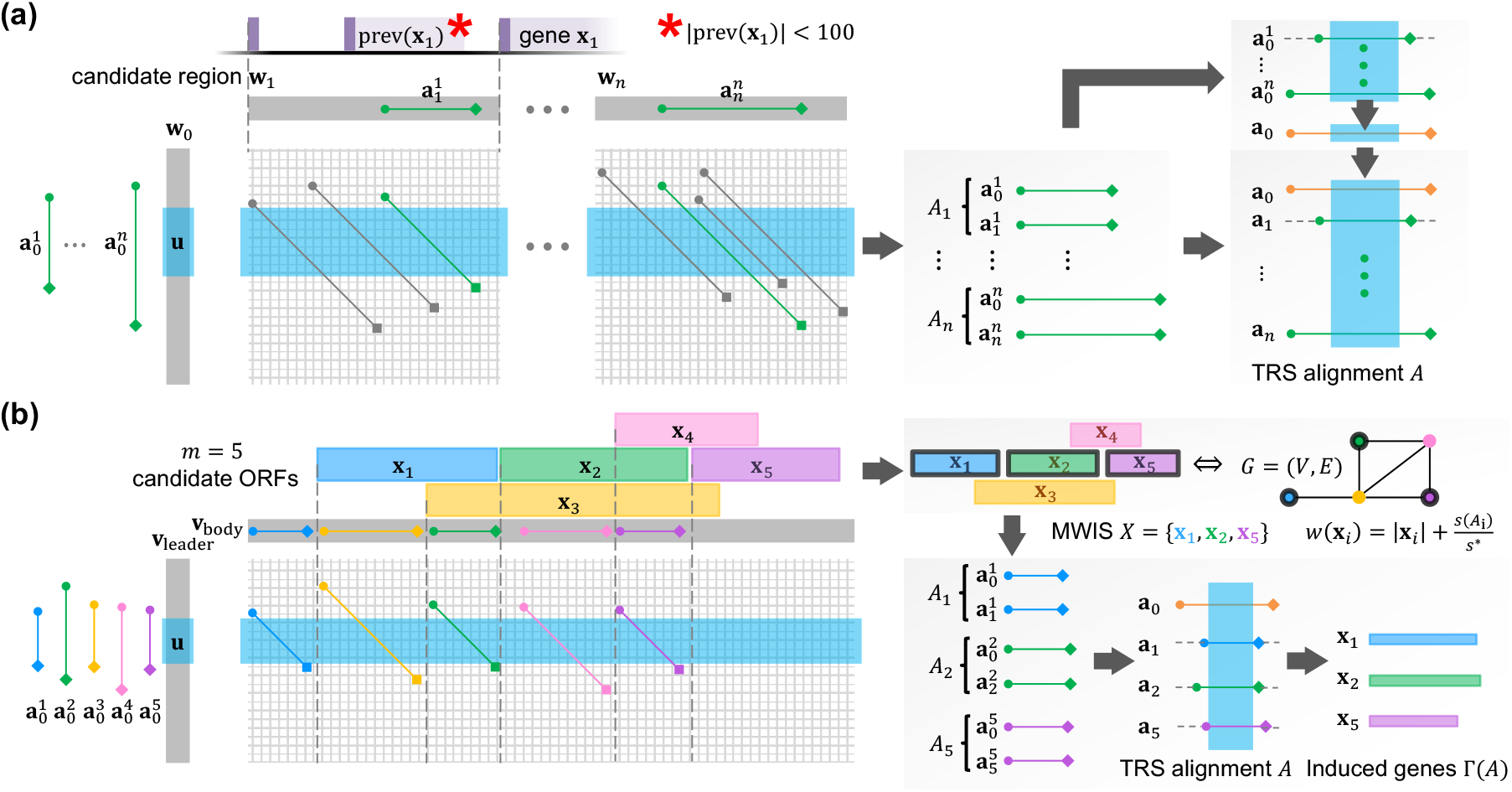
Algorithm details. (a) Given genes **x**_0_, *…*, **x**_*n*_, we obtain candidate regions **w**_0_, *…*, **w**_*n*_ by identifying upstream ORFs, skipping over ORFs if they are of length less than 100 nt (indicated by ‘*’). CORSID-A solves the TRS-ID problem by sliding a window **u** through **w**_0_, solving *n* independent pair-wise dynamic programming problems, which together yield the optimal TRS alignment *A* for window **u**. (b) To solve the TRS-Gene-ID problem, CORSID additionally solves a maximum-weight independent set problem [11] on an interval graph defined by the candidate ORFs to simultaneously identify an optimal pair (*A*, Γ(*A*)) for window **u**.

### 3.2 Solving the TRS AND GENE IDENTIFICATION problem

In the TRS-Gene-ID problem, we require two sequences: **v**_leader_ which contains TRS-L **a**_0_ and **v**_body_ which contains each TRS-B **a**_1_, *…*, **a**_*n*_. We describe a heuristic to partition a genome **v** into **v**_leader_ and **v**_body_ in Appendix A.3. This heuristic is performed in *O*(*m*^2^) time where *m* is the number of ORFs in **v**.

We will now define the relationship between a TRS alignment *A* = [**a**_0_, *…*, **a**_*n*_]^*T*^ and the set Γ(*A*) of induced genes. Upon removing (flanking) gaps, each aligned sequence **a**_*i*_ corresponds to a contiguous subsequence **v**_*i*_ of the viral genome **v**. Specifically, **v**_0_ occurs in **v**_leader_ and **v**_*i*_ occurs in **v**_body_ (where *i* ≥ 1). By Definition 4, each subsequence **v**_*i*_ has positions that are aligned with the core sequence **c**(*A*).

These aligned positions induce the subsequence 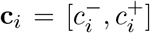 of length equal to **c**(*A*). Note that while **c**_0_ = **c**(*A*), it may be that **c**_*i*_ *≠* **c**(*A*) where *i* ≥ 1 due to mismatches. Importantly, there are coronaviruses where the last three nucleotides of the core sequence within a TRS-B coincide with the start codon of the associated gene (Fig. S1). As such, we have the following definition.

#### Definition 9.

Let *A* = [**a**_0_, *…*, **a**_*n*_]^*T*^ be a TRS alignment and let 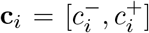 be the subsequence of **a**_*i*_ that is aligned to the core sequence **c**(*A*). The ORF *associated* with TRS-B **a**_*i*_ is the unique ORF **x** where position 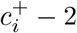 occurs within the candidate region of **x**.

As discussed, there may not exist an ORF associated with a TRS-B **a**_*i*_, which may happen when the TRS-B is located near the 3’ end of the genome. Given a TRS alignment *A* = [**a**_0_, *…*, **a**_*n*_]^*T*^, the set Γ(*A*) of induced genes equals the set of ORFs that are associated with **a**_1_, *…*, **a**_*n*_.

To solve the TRS-Gene-ID problem, we take a similar sliding window approach that we used to solve the TRS-ID problem. That is, we consider all subsequences **u** within **v**_leader_ of length *ω* and solve a constrained version of the TRS-Gene-ID problem, additionally requiring that the sought TRS alignment *A* has a core sequence **c**(*A*) that fully contains **u**, using the following two steps. First, we construct a DP table similar to the previous table used in TRS-ID problem in *O*(|**v**_leader_||**v**_body_|) time, and for each ORF, we select the alignment with the highest score in the corresponding candidate region. Second, given these ORFs and corresponding alignments, we build a vertex-weighted interval graph combining ORF lengths and alignment scores as weights. To identify the optimal TRS alignment *A* and associated genes Γ(*A*), we solve a maximum-weight independent set (MWIS) on this graph in *O*(*m*) time, where *m* is the number of candidate ORFs in **v**_body_ (Appendix A.4 and Fig. 2b). Each instance of the constrained TRS-Gene-ID problem takes *O*(|**v**_leader_||**v**_body_|+ *m*) time. Since the number of windows of length *ω* in **v**_leader_ is *O*(|**v**_leader_|), the total running time of CORSID to solve the TRS-Gene-ID problem is *O*(|**v**_leader_|^2^|**v**_body_ + **v**_leader_|*m*). In practice, the number *m* of candidate ORFs in **v**_body_ ranges from 21-92, the length **v**_leader_ of leader region ranges from 171-716 and the length |**v**_body_|of the body region ranges from 6280-11462 across all the coronaviruses studied in this paper. Finally, to obtain biologically meaningful solutions, we employ a progressive approach and consider overlapping genes (see Appendix A.5 for details).

## 4 Results

To evaluate the performance of CORSID-A and CORSID, we downloaded the same set of 505 assembled coronavirus genomes previously analyzed by SuPER [28] from GenBank along with their annotation GFF files, indicating gene locations. To benchmark methods for the TRS-ID problem, we assessed each method’s ability to correctly identify TRS-L as well as identify a TRS-B upstream of each gene. For the TRS-Gene-ID problem, we additionally assessed each method’s ability to identify ground-truth genes. Appendix B.1 describes how we established the set of genes and locations of TRS sites in the coronavirus genomes. We excluded 35 genomes due to incomplete leader sequences, thus lacking TRS-L. We excluded two more genomes due to empty GFF files, thus lacking gene annotations. The remaining 468 genomes comprised all four genera of the *Coronaviridae* family and spanned a total of 22 subgenera (Table S2).

### 4.1 CORSID-A finds TRS-L and TRS-Bs with higher accuracy than existing methods

We begin by comparing the performance of CORSID-A with MEME and SuPER for the TRS Identifi-cation problem. Recall that MEME is a general-purpose motif detection algorithm [2], whereas SuPER is specifically designed for identifying core sequences within coronavirus genomes annotated with genes [28]. To run CORSID-A, we extracted candidate regions **w**_1_, *…*, **w**_*n*_ upstream of annotated genes **x**_0_, *…*, **x**_*n*_ as described in Definition 11. The minimum length *ω* of core sequence is set to 7 following existing literature [8], and we use a minimum alignment score of *τ* = 2. We provided MEME with the same candidate regions **w**_0_, *…*, **w**_*n*_, and ran it in “zero or one occurrence per sequence” mode. As for SuPER, we analyzed the previously reported results on the same 468 sequences considered here. Detailed commands and parameters can be found in Appendix B.2.

As shown in Fig. 3b, CORSID-A correctly identified TRS-Ls in 466 out of 468 genomes, reaching a higher accuracy (99.6%) than MEME (442 genomes, 94.4%), but was outperformed by SuPER, which was correct in 467 genomes (99.8%). The two genomes where our method failed are outliers in their respective subgenera, indicative of possible sequencing errors (Appendix B.4). Fig. S2 discusses the one genome where SuPER failed to identify TRS-L correctly, showing that the TRS-L sequence identified by our method is supported by both secondary structure information as well as a split read in a corresponding RNA sequencing sample. *Split reads* map to non-contiguous regions of the viral genome and provide direct evidence of template switching at TRS sites.

**Figure 3:**
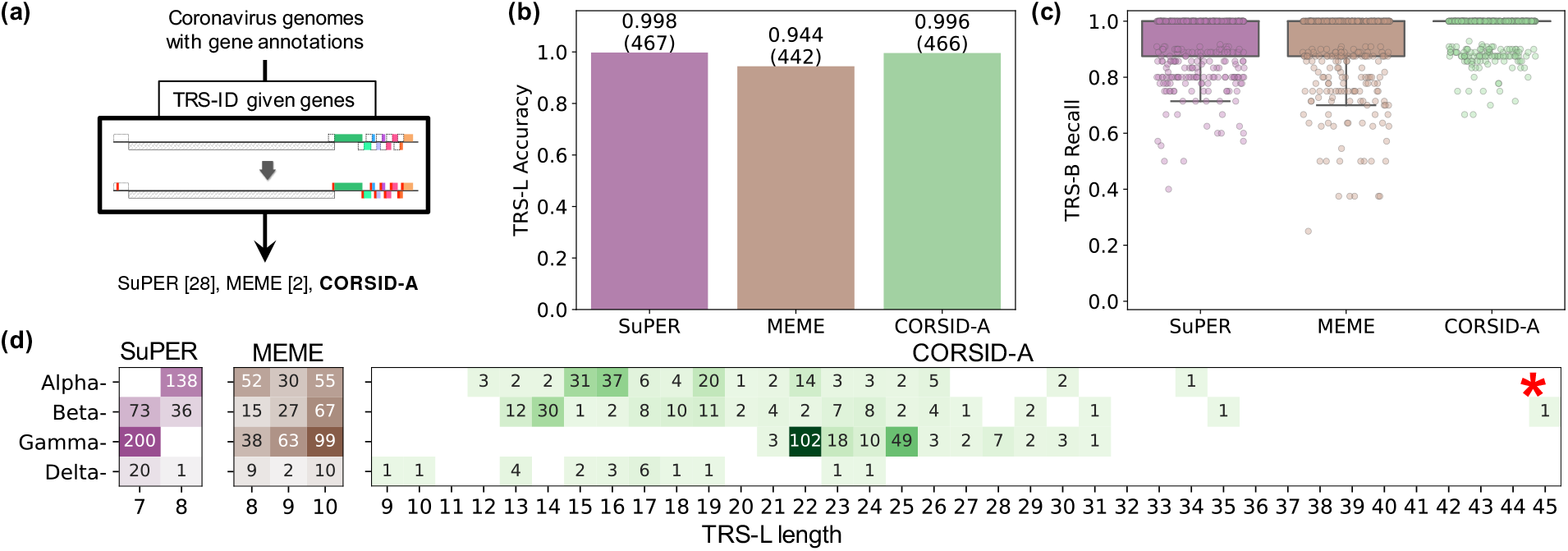
CORSID-A accurately identifies TRS-Ls and TRS-Bs. (a) We used SuPER [28], MEME [2] and CORSID-A to identify TRS sites in 468 coronavirus genome with known gene locations. (b) The fraction of genomes for which the three methods identified the TRS-L correctly. (c) The fraction of genes of the genomes for which the three methods identified the corresponding TRS-B site correctly. (d) Number of coronavirus genomes of the four genera of the *Coronaviridae* family with different lengths of the TRS-L identified by the three methods. Fig. S4 provides the TRS alignment identified by CORSID-A for the genome indicated by ‘*’.

Of note, SuPER uses additional information to identify TRS-L and TRS-B sites compared to MEME and CORSID-A. That is, SuPER requires the user to specify the genus of origin for each input sequence, which is used to obtain a genus-specific motif of the core sequence from a look-up table. This motif is used to identify matches along the genome. In addition, SuPER takes as input the 5’ UTR secondary structure, restricting the region in which the TRS-L occurs until the fourth stem loop (SL4). Importantly, while CORSID-A does not rely on any prespecified motif, taxonomic or secondary structure information, our method identified more TRS-Bs than either SuPER or MEME (Fig. 3c). Specifically, we define the *TRS-B recall* as the fraction of genes for which TRS-Bs were identified. While the median TRS-B recall of all three methods is 1, CORSID-A found putative TRS-Bs of all genes in 387 genomes (82.7%), while SuPER and MEME did so in only 290 (62.0%) and 315 (67.3%) genomes, respectively.

To validate the identified TRS sites, we examined split reads in publicly available RNA-sequencing data of cells infected by coronaviruses. Here we considered two samples, SRR1942956 and SRR1942957, of SARS-CoV-1-infected cells (NC_004718) with a median depth of 2940× and 2765×, respectively. The TRS-B region predicted by CORSID-A is supported by 246 reads in sample SRR1942956 and 233 reads in sample SRR1942957, whereas the TRS-B region predicted by SuPER is supported by only 1 read in each sample (Fig. S3a). Our method was able to identify these positions due to use of flanking positions rather than focusing on identifying a short 6− 7 nt motif as done by SuPER.

Recombination plays a crucial role in the evolution of RNA viruses, and results from homologous or non-homologous template switching. In particular for coronaviruses, template switching occurs at TRS sites during discontinuous transcription [9], making these sites prone to recombination events. CORSID-A uses local alignment to identify TRSs, and unlike SuPER, is not restricted to identifying regulatory sequences of a fixed length. Therefore, as a by-product, our method will be able to find evidence for homologous recombination at these sites. Specifically, even though the length of the core sequence is fixed at 7, the length of the TRS-Ls identified by our method ranges from 9 to 45 (median: 22). This corroborates previous findings showing that recombination hotspots in coronaviruses are colocated with TRS sites [28]. In particular, the longest TRS-B with a length of 45 nucleotides occurs upstream of gene *ns12.9* of NC 006213 with only 6 mismatches showing strong evidence of recombination (Fig. S4). In contrast, the core sequence identified by SuPER and MEME (Fig. 3d and Fig. S5) are at most 10 nt long. Furthermore, we note there is experimental evidence that not only the core sequence but also flanking nucleotides play an important role in discontinuous transcription [26]. This demonstrates the importance of identifying larger regulatory sequences, as done by our method, rather than identifying shorter recurring motifs as done by SuPER and MEME. In summary, CORSID-A outperforms existing methods, such as SuPER and MEME, in identifying TRS sites in coronavirus genomes with given gene locations.

### 4.2 CORSID identifies genes with higher accuracy than existing methods

We now focus on the TRS-Gene-ID problem, where we compared CORSID to two general-purpose gene finding methods: Glimmer3 [5, 21] and Prodigal [12, 13]. Each method was given as input the complete, unannotated genome sequence of each of the 468 coronaviruses. Following recommended instructions, we ran Glimmer3 by first building the required interpolated context model (ICM) on each genome sequence separately. We ran Prodigal in meta-genomics mode. For CORSID, we used window length *ω* = 7 and progressively reduced the score threshold *τ* from 7 to 2. We refer the reader to Appendix B.2 for the precise commands used to run previous tools, and to Appendix B.3 for details on how the predicted set of genes are compared to the ground truth.

Fig. 4a shows that CORSID outperformed Glimmer3 and Prodigal in terms of both precision and recall. The median precision and recall of CORSID is 0.818 and 1.00, respectively, whereas the median precision and recall is 0.556 and 0.556, respectively, for Glimmer3, and 0.636 and 0.667, respectively, for Prodigal. Additionally, the first quartile of both precision and recall for CORSID is greater than the third quartile of these metrics for either Glimmer3 and Prodigal, showing a significant performance advantage for CORSID. While Prodigal and Glimmer3 do not have the capability to identify TRS sites, CORSID identifies these regulatory sites in addition to the genes. Specifically, compared to CORSID-A, which identified TRS-L correctly for 466 (99.6%) genomes, CORSID does so for 443 (94.7%) genomes (Fig. S6). This is a modest reduction in performance, especially when taking into account that CORSID, unlike CORSID-A, is not given any additional information apart from the complete, unannotated genome sequence. Analyzing the previously discussed SARS-CoV-1 genome (NC 004718), we found that CORSID identified the same 10 genes as CORSID-A, while Prodigal missed four genes and Glimmer3 missed two genes (Fig. S3b).

**Figure 4:**
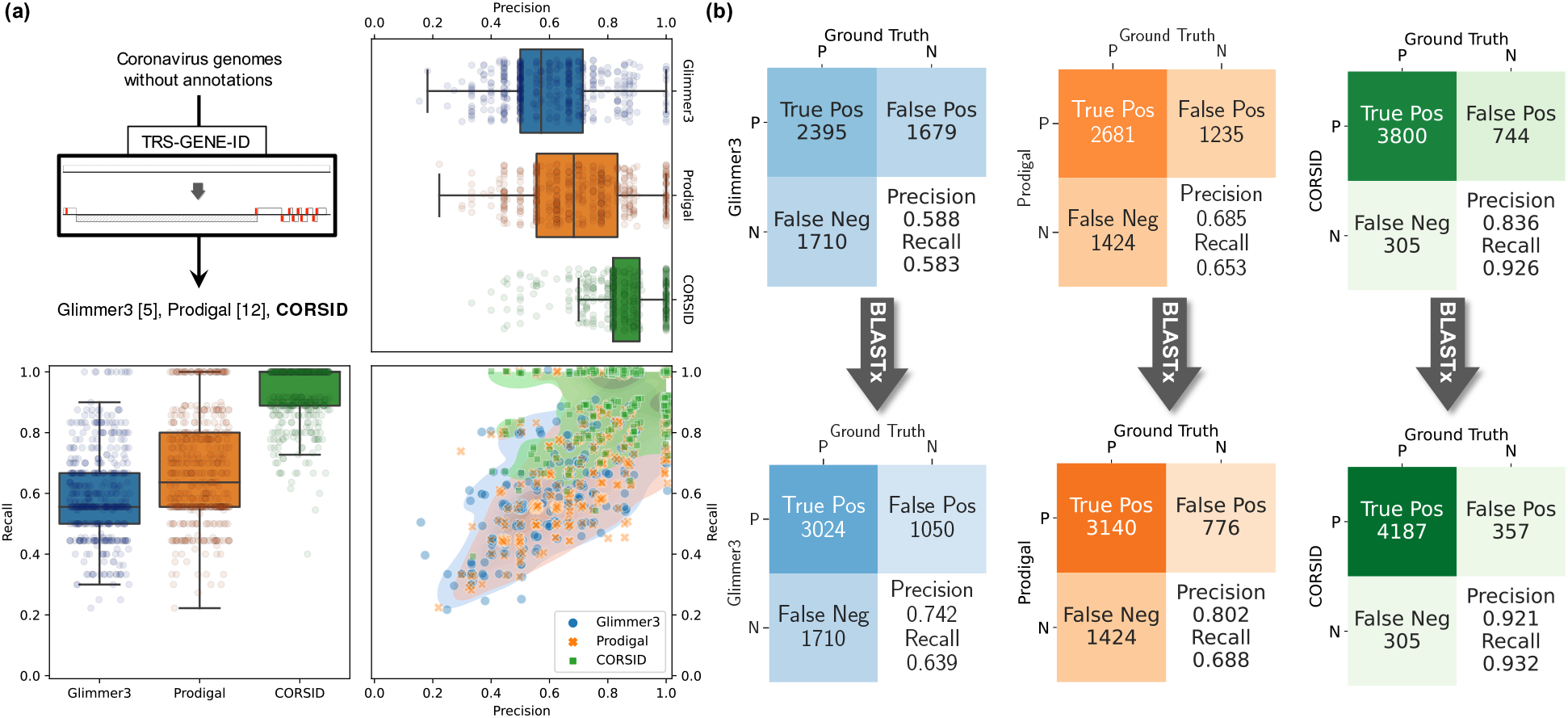
CORSID accurately identifies TRS-Ls, TRS-Bs, and genes. (a) Precision and recall of Glimmer3 [5], Prodigal [12], and CORSID for gene prediction in 468 genomes. For clarity, we added a small jitter (drawn from *N* (0, 2.5 × 10^*−*5^)) to the 2D distribution plot. (b) Confusion matrices of the ground truth genes and the predicted genes by the three methods. In order to account for poorly annotated genomes, we used BLASTx [1] to verify the false positive genes predicted by the three methods. The adjusted confusion matrices are shown in the row below.

Given the large discrepancy between precision and recall for CORSID, we investigated whether genomes were poorly annotated, leading to incorrect false positives. To that end, we used BLASTx [1] to transfer gene annotation from well-annotated genomes to poorly-annotated genomes. To reduce computation time, we only assessed false positive (FP) genes occurring in **v**_body_. We reclassified a FP gene as a true positive (TP) if the alignment reported by BLASTx spans at least 95% of both the query sequence (the FP gene) and the hit sequence in the database (detailed in Appendix B.2). The resulting confusion matrices of the predicted and the ground-truth genes are shown in Fig. 4b. Although BLASTx found 629 and 459 matches in the database for FPs of Glimmer3 and Prodigal, respectively, which is more than 387 for CORSID, CORSID maintains its lead in absolute numbers for TP and FP. As such, CORSID achieved the highest pooled precision and recall both before and after the BLASTx verification. In summary, CORSID accurately identifies TRSs and genes given just the unannotated genome, outperforming existing gene finding methods.

### 4.3 CORSID enables *de novo* identification of TRS sites and genes

To demonstrate how users can use CORSID to annotate genes and identify TRS-L and TRS-Bs given a newly assembled genome, we analyzed a previously-excluded genome that lacks gene annotation (genome DQ288927). This genome is 27534 nt long, which we provided as input to CORSID, Glimmer3 and Prodigal. CORSID identified nine genes spanning 91.66% of the genome, all of which match annotated genes in other *Igacoviruses* sequences in the BLASTx database (Fig. 5). By contrast, Glimmer3 identified a total of six genes spanning 80.52% of the genome, five of which match genes in the BLASTx database. On the other hand, Prodigal found six genes, all of which were present in the database, spanning 84.22% of the genome. In summary, CORSID identified more genes than existing methods, all of which occurred in homologous previously-annotated genomes in the BLASTx database, demonstrating that CORSID can be used to accurately annotate coronavirus genomes.

**Figure 5:**
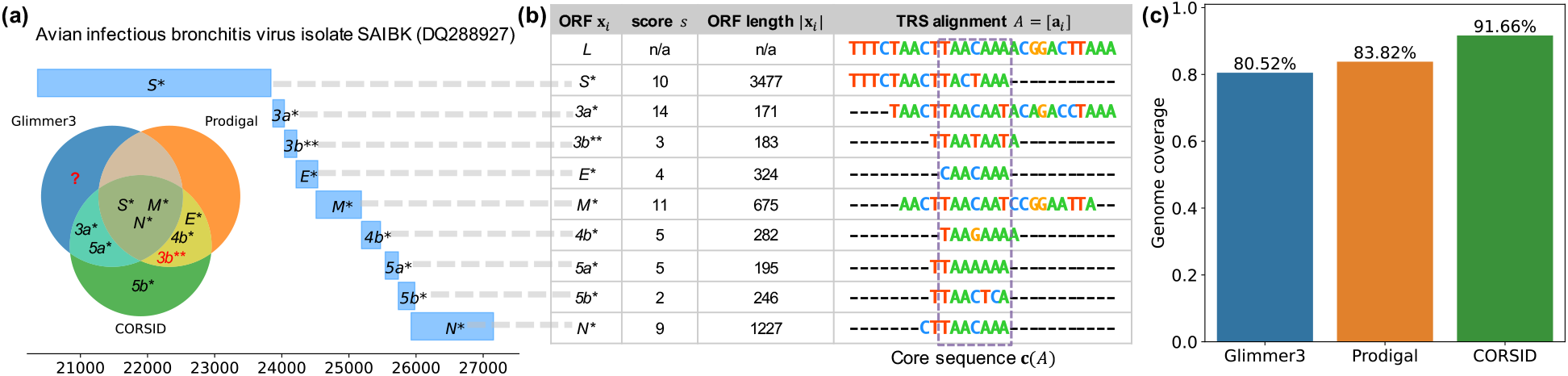
CORSID accurately finds genes in an unannotated *Igacovirus* genome (DQ288927). (a) The position of the genes identified by CORSID. The Venn diagram shows the genes found by CORSID, Glimmer3 and Prodigal. “*” indicates ≥ 95% query/hit coverage, and “**” indicates a hit coverage of 93.5% and a query coverage of 98.3%. (b) TRS alignment for genes identified by CORSID. (c) The fraction of positions in **v**_body_ covered by genes identified by the three methods.

## 5 Discussion

In this paper, we investigated the hypothesis that the presence of transcription regulatory sequences in coronavirus genomes can be leveraged to simultaneously infer these regulatory sequences and their associated genes in a synergistic manner. To that end, we formulated the TRS Identification (TRS-ID) problem of identifying TRS sites in a coronavirus genome with given gene locations, and the general problem, the TRS and Gene Identification (TRS-Gene-ID) problem of simultaneous identification of genes and TRS sites given only the coronavirus genome. Underpinning both problems is the notion of a TRS alignment, which extends the previous concept of core sequences to include flanking nucleotides that provide additional signal. Our proposed method for the first problem, CORSID-A, is based upon a dynamic programming formulation which extends the classical Smith-Waterman recurrence [24]. CORSID, which solves the general problem, additionally incorporates a maximum-weight independent set formulation on an interval graph to identify TRS sites and genes.

Using extensive experiments on 468 coronavirus genomes, we showed that CORSID-A outperformed two motif-based approaches, MEME [2] and SuPER [28]. Additionally, we showed that CORSID outperformed two general-purpose gene finding algorithms, Glimmer3 [5, 21] and Prodigal [12]. We performed direct validation of TRS sites predicted for the SARS-CoV-1 genome (NC 004718), showing that the TRS sites identified by our method are more strongly supported by split reads in RNA-seq samples than the TRS sites identified by SuPER. Lastly, we demonstrated that CORSID enables *de novo* identification of TRSs and genes in newly assembled coronavirus genomes by applying it on a previously unannotated coronavirus (DQ288927) belonging to the *Igacovirus* subgenus.

There are several avenues for future research. First, CORSID currently requires the complete genome as input to identify the TRS sites and the genes. We plan to extend our method to allow gene identification in the several coronaviruses available in GenBank with only partial reference genomes by leveraging knowledge from other coronaviruses with complete genomes with similar TRS sites. Second, while in this study we only focused on coronaviruses, discontinuous transcription occurs in all viruses in the taxonomic order of *Nidovirales*. However, CORSID, which assumes a single TRS-L region in the genome, cannot be directly applied to other families of viruses within *Nidovirales* such as the family *Mesoniviridae* that contain multiple TRS-L regions in the genome. Incorporating such features and extending CORSID to all *Nidovirales* viruses is a useful direction of future work. Finally, currently CORSID requires the reference genome of the virus as input. In the future, we plan to extend this method to simultaneously discover the reference genome and the core-sequences of the virus using *de novo* assembly. We envision that this will enable simultaneous genome assembly and gene annotation of coronaviruses.

## Acknowledgements

This material is based upon work supported by the National Science Foundation under award numbers CCF-1850502, CCF-2027669 and CCF-2046488. We thank Ayesha Kazi, Michael Xiang, and Yichi Zhang for developing the web-based visualization tool of CORSID solutions.

## A Supplementary Methods

### A.1 Obtaining candidate regions for the TRS-ID problem

Intuitively, a candidate region for gene **x**_*i*_ corresponds to the region 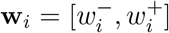 composed of positions *w*^−^ ≤ *p* ≤ *w*^+^ such that any sgmRNA starting at *p* will lead to the translation of ORF **x**_*i*_ by the ribosome. Note that the first ORF **x**_0_ corresponds to *ORF1ab*. Accordingly, we restrict 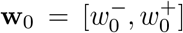 to match exactly the leader region, spanning the start of the genome at the 5’ end until the start codon of **x**_0_, *i.e* 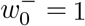 and 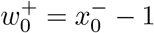. To define the remaining candidate regions **w**_1_, *…*, **w**_*n*_, we must take ribosomal leaky scanning into account, where the ribosome does not initiate translation at the first ‘ATG’ it encounters [14]. To model this, we make use of the fact that coronovirus genes have a length of at least 100 nucleotides. Specifically, when determining the candidate region of a gene, we skip over a previous ORF in case its length is less than 100. To that end, we introduce the following function.

#### Definition 10.

Function prev(*p*) returns the first ORF **x** = [*x*^−^, *x*^+^] upstream of position *p* in the genome, *i.e*. for ORF **x** returned by prev(*p*) it holds that *x*^−^ *< p* and there exists no ORF **y** = [*y*^−^, *y*^+^] such that *x*^−^ *< y*^−^ *< p*. If no such ORF **x** exists then prev(*p*) = [0, 0]. Moreover, prev(0) = [0, 0].

Using this function, we define a TRS-B candidate region **w** of an ORF **x** as follows.

#### Definition 11.

Let **x** = [*x*^−^, *x*^+^] be an ORF, and let **y** = [*y*^−^, *y*^+^] = prev(*x*^−^) and **z** = [*z*^−^, *z*^+^] = prev(*y*^−^) be the previous two ORFs. The *candidate region* **w** = [*w*^−^, *w*^+^] of ORF **x** ends at the start of **x**, *i.e. w*^+^ = *x*^−^ − 1, and begins at the first position of the genome if **x** has no previous ORF or the only preceding ORF **y** has a length smaller than 100; **w** begins at the first ORF **y** if the length of **y** is at least 100; otherwise **w** begins at the second ORF **z** if the first ORF **y** has a length smaller than 100 nucleotides. That is,

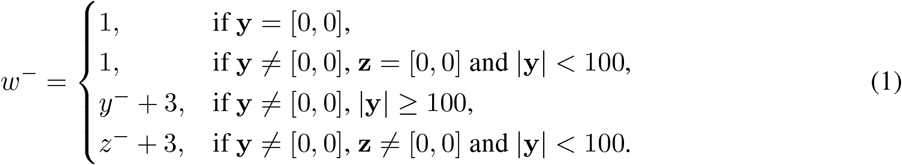

Finally, to remove overlap among candidate regions **w**_0_, *…*, **w**_*n*_, we set 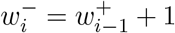 if 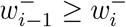 for all *i* ∈ {1, *…, n*}.

### A.2 Constrained TRS-ID problem

Here, we introduce the following constrained version of the TRS-ID problem.

**Problem 3** (Constrained TRS Identification (TRS-ID-**u**)). Given non-overlapping sequences **w**_0_, *…*, **w**_*n*_, and a subsequence **u** of **w**_0_, find a TRS alignment *A* = [**a**_0_, *…* **a**_*n*_]^*T*^ such that (i) **a**_*i*_ corresponds to a subsequence in **w**_*i*_ for all *i* ∈ {0, *…, n*}, (ii) **u** is a subsequence of the core sequence **c**(*A*), and (iii) the alignment has maximum score *s*(*A*).

Let us focus on solving a single TRS-ID-**u** problem instance, where we are given non-overlapping sequences **w**_0_, *…*, **w**_*n*_ and a subsequence **u** of **w**_0_. Each such instance decomposes into *n* TRS-ID-**u** instances each with exactly two sequences. That is, for any *i* ∈ 1, *…, n*, we seek a TRS alignment 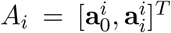 of sequences 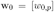 and 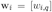 such that the induced core sequence **c**(*A*_*i*_) contains **u** = [*u*^−^, *u*^+^]. This is a variant of local alignment [24] with three key differences: (i) alignment *A*_*i*_ may not contain gaps, (ii) *A*_*i*_ must span *u*^−^ and (iii) *A*_*i*_ must span *u*^+^. Letting be ℓ the relative position of *u*^−^ in **w**_0_, we obtain

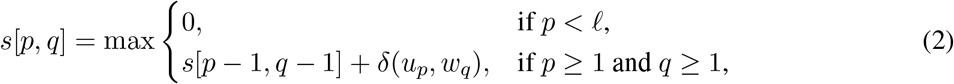

where *s*[*p, q*] indicates the optimal score of a constrained TRS alignment between *w*_0,1_ *… w*_0,*p*_ and *w*_*i*,1_ *… w*_*i,q*_. The desired solution is then as follows.

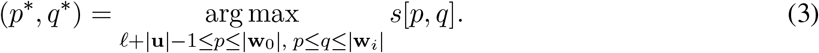

Note that the recurrence (2) lacks the two cases of the Smith-Waterman [24] recurrence corresponding to a gap (*i.e. s*[*p* − 1, *q*] and *s*[*p, q* − 1]), thus satisfying constraint (i). Constraints (ii) and (iii) are satisfied because the first case, corresponding to initiating the alignment, is only enabled when *p <* thus covering *u*^−^, and by (3), we have that the alignment contains *u*^+^. Similarly to local alignment, we can identify (*p*^*^, *q*^*^) by filling out the table *s*[0, 0], *…, s*[|**w**_0_|, |**w**_*i*_|] using dynamic programming, and reconstruct the TRS alignment 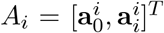 using a backtrace from (*p*^*^, *q*^*^). Letting 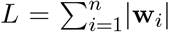 be the total length of candidate regions **w**_1_, *…*, **w**_*n*_, solving the *n* pairwise TRS-ID-**u** problems and obtaining the pairwise TRS alignments *A*_1_, *…, A*_*n*_ that cover **u** takes *O*(|**w**_0_ *L*|) time.

Given these pairwise alignments *A*_1_, *…, A*_*n*_ that each span **u**, we construct the final TRS alignment *A* = [**a**_0_, *…*, **a**_*n*_]^*T*^ as follows. First, we exclude alignments *A*_*i*_ that have a score less than the threshold *τ*. Second, the TRS-L sequence **a**_0_ equals the subsequence of **w**_0_ that spans the positions covered by all pairwise alignments, *i.e*. **a**_0_ spans exactly the positions of **w**_0_ covered by 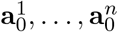. Third, we obtain the remaining gapped sequences **a**_1_, *…*, **a**_*n*_ of *A* by adding flanking gaps to each (ungapped) sequence 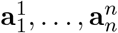 so as to match the unaligned letters of **a**_0_ (Fig. 2a). As flanking gaps do not incur a penalty, this operation will not change the total score, *i.e*. 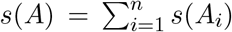. The running time of computing alignments *A*_1_, *…, A*_*n*_ and then subsequently merging them into *A* is dominated by the first step.

### A.3 Partitioning the genome into v_leader_ and v_body_ for the TRS-GENE-ID problem

To obtain these two sequences, **v**_leader_ and **v**_body_, for a given coronavirus genome, we developed a heuristic for identifying *ORF1ab*, the largest gene in coronavirus genomes. This heuristic begins by enumerating all ORFs **x**_1_, *…*, **x**_*m*_ in the genome (Definition 5). As *ORF1ab* is the result of a frameshift upstream of the stop codon of *ORF1a* [18], we extend each enumerated ORF **x**_*i*_ by performing either a − 1 or − 2 frameshift and subsequently scanning for an in-frame stop codon. We select the frameshift that results in the largest extended ORF, obtaining extended ORFs **y**_1_, *…*, **y**_*m*_. We designate the largest ORF among this set as *ORF1ab*. Finally, we set **v**_leader_ as the region from the start of the genome until the 5’ coordinate of *ORF1ab*. As the TRS-B of the first gene downstream of *ORF1ab* may reside within *ORF1ab*, we set **v**_body_ as the region starting from 200 nucleotides upstream of the 3’ coordinate of *ORF1ab* until the 3’ end of the genome.

### A.4 Constrained TRS-GENE-ID problem

Here, we introduce the constrained version of the TRS-Gene-ID problem.

**Problem 4** (Constrained TRS and Gene Identification (TRS-Gene-ID-**u**)). Given leader region **v**_leader_, body region **v**_body_ and a subsequence **u** of **v**_leader_, find a TRS alignment *A* = [**a**_*i*_] such that (i) **a**_0_ corresponds to a subsequence in **v**_leader_, (ii) **a**_*i*_ corresponds to a subsequence in **v**_body_ for all *i* ≥ 1, (iii) **u** is a subsequence of the core sequence **c**(*A*), (iv) *A* is concordant, and (v) *A* induces the set Γ(*A*) of genes with maximum genome coverage *g*(*A*) and subsequently has maximum score *s*(*A*).

We solve this problem in two steps. First, we use dynamic programming to compute *s*[*p, q*] for all values of 0 ≤ *p* ≤ |**v**_leader_| and 0 ≤ *q* ≤ |**v**_body_|, *i.e*. the optimal score *s*[*p, q*] of a TRS alignment between *v*_leader,1_ *… v*_leader,*p*_ and *v*_body,1_ *… v*_body,*q*_ constrained to contain **u**. The quantity *s*[*p, q*] is defined using the same recurrence as for the TRS-ID-**u** problem – Eq. (2) – and the complete table can be filled out using dynamic programming in *O*(| **v**_leader_||**v**_body_|) time.

Second, let **x**_1_, *…*, **x**_*m*_ be the candidate ORFs in **v**_body_, each with a length of at least 100 nucleotides (Fig. S7). For each ORF **x**_*i*_ we find the position (*p, q*) that encodes the maximum scoring alignment 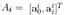 where 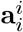 is associated with **x**_*i*_. We remove ORFs **x**_*i*_ whose maximum scoring alignment *A*_*i*_ has a score *s*(*A*_*i*_) less than the user-specified score threshold *τ*. Let *s*^*^ indicate the maximum score among all TRS alignments *A*_1_, *…, A*_*m*_. Then, we construct a vertex-weighted interval graph *G* = (*V, E*) whose vertices *V* correspond to the intervals 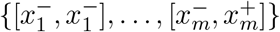 of the candidate ORFs. There is an edge (**x**_*i*_, **x**_*j*_) if and only if the two corresponding intervals overlap, *i.e*. 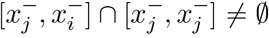. To capture the lexicographical ordering of the objective functions, *i.e*. first the genome coverage and then the score, each vertex/interval **x**_*i*_ is assigned weight

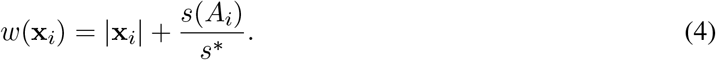

In other words, among ORFs with the same length, we prefer those that have an associated TRS alignment with largest score. Finally, we solve a maximum-weight independent set (MWIS) problem, which can be done in *O*(|*V* |) time for interval graphs [11]. The maximum-weight independent set *X* = {**x**_*π*(1)_, *…*, **x**_*π*(|*X*|)_} directly corresponds to the induced genes Γ(*A*) of the TRS alignment *A* that can be constructed by merging pairwise TRS alignments *A*_*π*(1)_, *…, A*_*π*(|*X*|)_ following the same procedure described in Section 3.1. For each window **u** of fixed length *ω*, the first step takes *O*(|**v**_leader_||**v**_body_|) time and the second step takes *O*(|**v**_leader_||**v**_body_| + *m*) time.

### A.5 Practical considerations to solve the TRS-GENE-ID problem

In this section we discuss practical considerations for identifying genes in coronaviruses and the way they are addressed in CORSID.

#### Overlapping genes

In practice, coronavirus genes may overlap. That is, the start codon of a gene can be located within another gene. To support such cases, we shrink candidate ORFs **x** = [*x*^−^, *x*^+^] by 5% prior to solving the MWIS problem, *i.e*. the interval graph *G* = (*V, E*) has shortened intervals **x**^i^ = [*x*^−^ +*α, x*^+^ −*α*] where *α* = 0.05|**x**|.

#### Progressive approach

To obtain biologically meaningful solutions, we solve the TRS-Gene-ID problem in a progressive manner. More specifically, given user-specified parameters (*τ*_min_, *τ*_max_), we start with setting *τ* = *τ*_max_ and solve the problem for a fixed window **u**. Then, for every subsequent iteration we decrement *τ* and require the solution to contain all ORFs that were identified in the previous iteration. The final iteration occurs when *τ* = *τ*_min_, yielding the final solution for window **u**. We consider all sliding windows **u** within **v**_leader_ and return the solution with maximum genome coverage and subsequently maximum score. We show that the progressive approach performs better than solving the TRS-Gene-ID problem directly using *τ* = *τ*_min_ or *τ* = *τ*_max_ in Fig. S8.

## B Supplementary Results

### B.1 Establishing the ground truth of gene and TRS locations in coronavirus genomes

We established a ground-truth set of genes by processing the GFF annotation files and extracting a set of genes for each genome, removing duplicates and incomplete ORFs. In particular, we removed 10 ORFs (8 annotated as *N* and 2 annotated as ‘unknown’) that had duplicate names and were completely covered by another gene with the same name. This resulted in a median number of 8 genes per genome (min 3 and max 14, Fig. S9). We excluded two genomes that had no genes in their annotation (DQ288927 and EU526388). We used the following approach to establish the ground truth for TRS-B sites. As discussed in Section 3.1, we identified *candidate regions* (Definition 11) for each gene. If a method identified a TRS-B contained within any of the candidate regions, then we counted this as the method recalling the TRS-B for the corresponding gene.

We established ground-truth locations of TRS-Ls using the fact that these regulatory sequences occur between the second (SL2) and fourth stem loop (SL4) in the 5’ untranslated region (UTR) [17]. For *Sarbecovirus* genomes, a subgenus of the *Betacoronavirus* genus, we required TRS-Ls to occur in stem loop (SL3) [4]. Specifically, we analyzed leader regions upstream of *ORF1ab*, its location taken from the GFF annotation or determined by our own heuristic in case this gene was absent, and performed a multiple sequence alignment using ClustalW2 [16] of all sequences within each subgenus. We then superimposed secondary structure information from Rfam [10] onto each alignment to identify the relevant stem loops in each viral sequence, obtaining for each sequence a small range in which TRS-L may occur (Fig. S10).

### B.2 Command-line arguments

#### MEME

We ran MEME v5.3.0 in mode “zero or one occurrence per sequence” (zoops) and maximum width of 10.

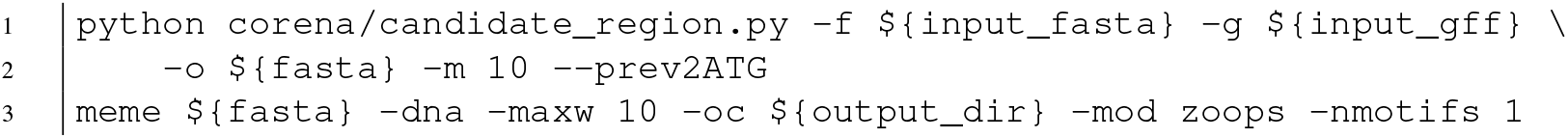

#### Glimmer3

We followed the steps written in the script `g3-from-scratch.csh` provided in the Glimmer3 package.

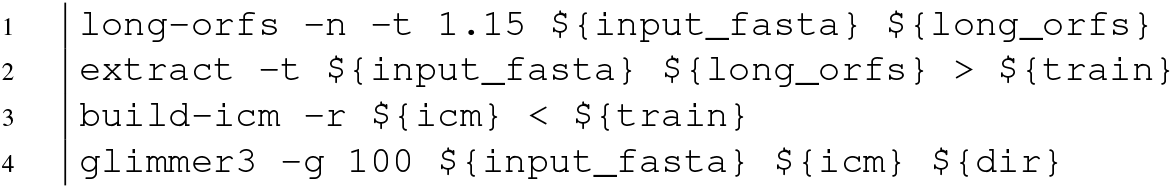

#### Prodigal

We ran Prodigal v2.6.3 in metagenomic mode.

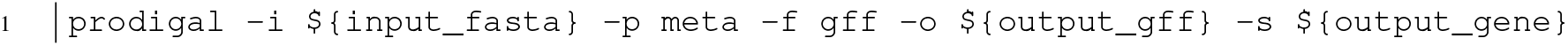

#### ClustalW2

We used ClustalW2 v2.1 to align sequences.

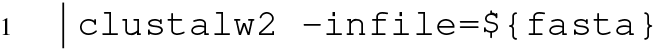

#### BLASTx

We ran BLASTx on a FASTA file containing FP genes with the following parameters. We used a subset of protein sequences from the official “nr” database (downloaded on October 7, 2021), containing all species under the taxonomic unit *Coronaviridae* (taxid:11118). From the BLASTx output, we extracted the top hit for each FP gene, and calculated the coverage between the aligned subsequence, and the query and hit sequences. In particular, the *hit coverage* is fraction of position in the hit sequence that are aligned. On the other hand, the *query coverage* is the fraction the query sequence that are aligned. We set the threshold to 95% for both hit and query coverage, restricting solutions to almost exact matches.

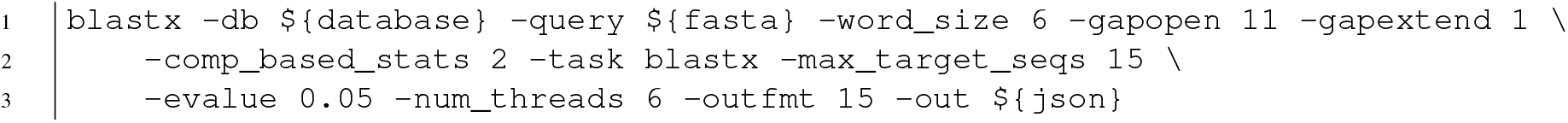

### B.2 Evaluating gene finding methods

To assess performance of gene finding methods, we say that a predicted gene **x** = [*x*^−^, *x*^+^] is correct provided there exists a ground-truth gene **y** = [*y*^−^, *y*^+^] in the same genome such that |*x*^−^ − *y*^−^|≤ 3 and |*x*^+^ − *y*^+^| ≤ 3. In other words, the start and end positions may be off by the length of at most one codon, accounting for variation in annotation (e.g. sometimes the stop codon is omitted from ORFs). Moreover, as *ORF1ab* is not a real ORF, i.e. the corresponding polypeptide 1ab results from a−1 frameshift, we treat this gene differently. That is, we say that a method correctly identified **y** = [*y*^−^, *y*^+^] = *ORF1ab* if it found two genes 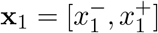 and 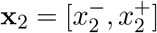 such that the start position of **x**_1_ is at most 3 nucleotides away from *y*^−^ and the stop position of **x**_2_ is at most 3 nucleotides away from *y*^+^. Note that CORSID will identify a single gene **x** matching *ORF1ab*, in which case **x** = **x**_1_ = **x**_2_. Using this definition we classify each predicted and ground-truth gene as either a *true positive* (TP), i.e. the predicted gene matches a ground-truth gene; *false positive* (FP), i.e. the predicted gene does not match any ground-truth gene); or *false negative* (FN), i.e. the ground-truth gene has no matching predicted gene.

### B.4 Two anomalous coronavirus genomes

CORSID-A was unsuccessful in identifying the correct TRS-L site in only two coronaviruses: MK211372 (from subgenus *Pedacovirus* of genus *Alphacoronovirus*) and MK472070 (unclassified subgenus of genus *Alphacoronovirus*).

To understand why CORSID-A failed on MK211372, we performed a multiple sequence alignment of the leader regions of all 45 genomes in the *Pedacovirus* subgenus. Inspecting the alignment, we see that MK211372 is an outlier, with multiple insertion/deletions in the TRS-L region compared to the other sequences (Fig. S11). This explains why CORSID-A was unable to accurately identify the TRS-L and TRS-B sites for this genome.

Since genome MK472070 has a known genus but unknown subgenus, we only aligned it to the covariance model of the alphacoronaviruses. From the alignment result we found a poor alignment in the TRS-L region. Based on the alignment, it resembles some nyctacoviruses, but it is still an outlier, as shown in Fig. S12. Specifically, the TRS-L consensus sequence, 5’-TCAACTAAAC-3’, differed significantly from the subsequence 5’-ACAATCTAAT-3’ of MK472070, with a Hamming distance of 5. Moreover, MEME also failed to identify TRS-L in this genome, and SuPER assigned a low confidence score to the identified TRS-Bs for important genes such as *S* and *N*.

In summary, we believe further investigation of genomes MK211372 and MK472070 is warranted in order to determine whether they harbor a TRS-L region or whether the deposited genome sequences are incomplete/incorrect.

### C Supplementary Figures and Tables

We have the following supplementary figures and tables.

- Fig. S1 shows an example of TRS regions overlapping with the start codon of the corresponding genes in coronaviruses.
- Fig. S2 shows CORSID finds the correct TRS-L in a *Sarbecovirus*, verified using a RNA-seq dataset.
- Fig. S3 shows RNA-seq data supports the TRS-B found by CORSID-A in SARS-CoV-1 (NC 004718), and CORSID identifies more genes than Glimmer3 and Prodigal in the same genome.
- Fig. S4 shows the TRS alignment in genome NC 006213 where CORSID find the longest alignment between TRS-L and TRS-B.
- Fig. S5 shows the number of coronaviruses with varying lengths of TRS-L regions broken down by the genus and the subgenus. Several coronaviruses of each genus have TRS-L regions much longer than the core sequences indicating recombination events.
- Fig. S6 shows that CORSID shows modest reduction in TRS-L accuracy compared to CORSID-A.
- Fig. S7 shows the distribution of length of annotated genes. Only 10 out of 3637 genes are shorther than 100 nt.
- Fig. S8 shows that CORSID achieves better performance when using the progressive approach rather than directly solving *τ* = *τ*_max_ = 7 or *τ* = *τ*_min_ = 2.
- Fig. S9 shows the histogram of length of annotated genes from 468 genomes, of which only 10 genes are shorter than 100 nt.
- Fig. S10 shows the distribution of the length of the TRS-L regions, separated by four genera.
- Fig. S11 shows the MSA of leader regions of pedacoviruses with genome MK211372 highlighted, indicating multiple INDELs.
- In Table S1, we compare features of CORSID, CORSID-A, and other methods. We show that CORSID is the first method for simultaneous identification of TRS sites and genes in coronaviruses.
- Table S2 shows the number of included and excluded genomes grouped by genera and subgenera.

**Table S1:** CORSID is the first method for simultaneous identification of TRS sites and genes in coronaviruses. This table shows the features of CORSID and CORSID-A along with three existing methods: MEME [2], Glimmer3 [5] and Prodigal [12].

|  | TRS-L identification | TRS-B identification | Gene identification |
| --- | --- | --- | --- |
| CORSID | ✓ | ✓ | ✓ |
| CORSID-A | ✓ | ✓ | ✗ |
| SuPER [28] | ✓ | ✓ | ✗ |
| MEME [2] | ✓ | ✓ | ✗ |
| Glimmer3 [5] | ✗ | ✗ | ✓ |
| Prodigal [12] | ✗ | ✗ | ✓ |

**Figure S1:**
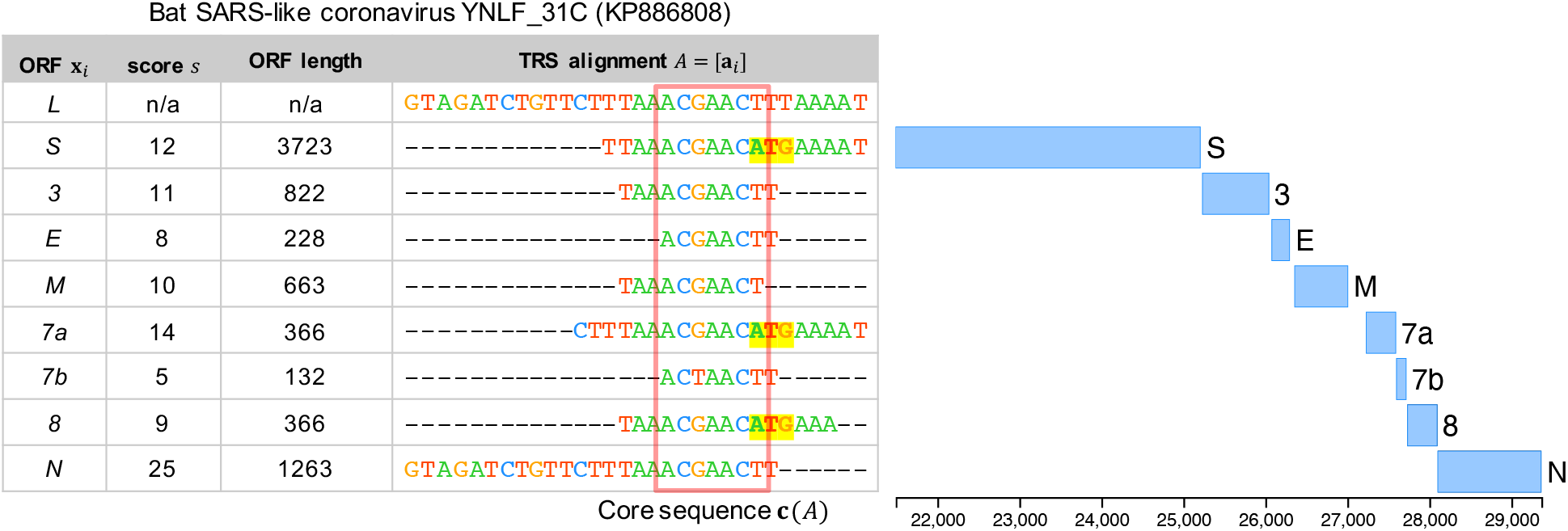
TRS sites may overlap with the start codon of the corresponding genes in coronaviruses. The TRS alignment identified by CORSID when applied to a *Sarbecovirus* genome KP886808, showing th start codons of gene *S, 7a*, and *8* are partially contained in core sequences (highlighted in yellow. We note that MEME identified the same TRS-L as our method. As a side note, the TRS-B of gene *N* matches 25 nucleotides of the TRS-L, indicative of a possible a recombination event.

**Figure S2:**
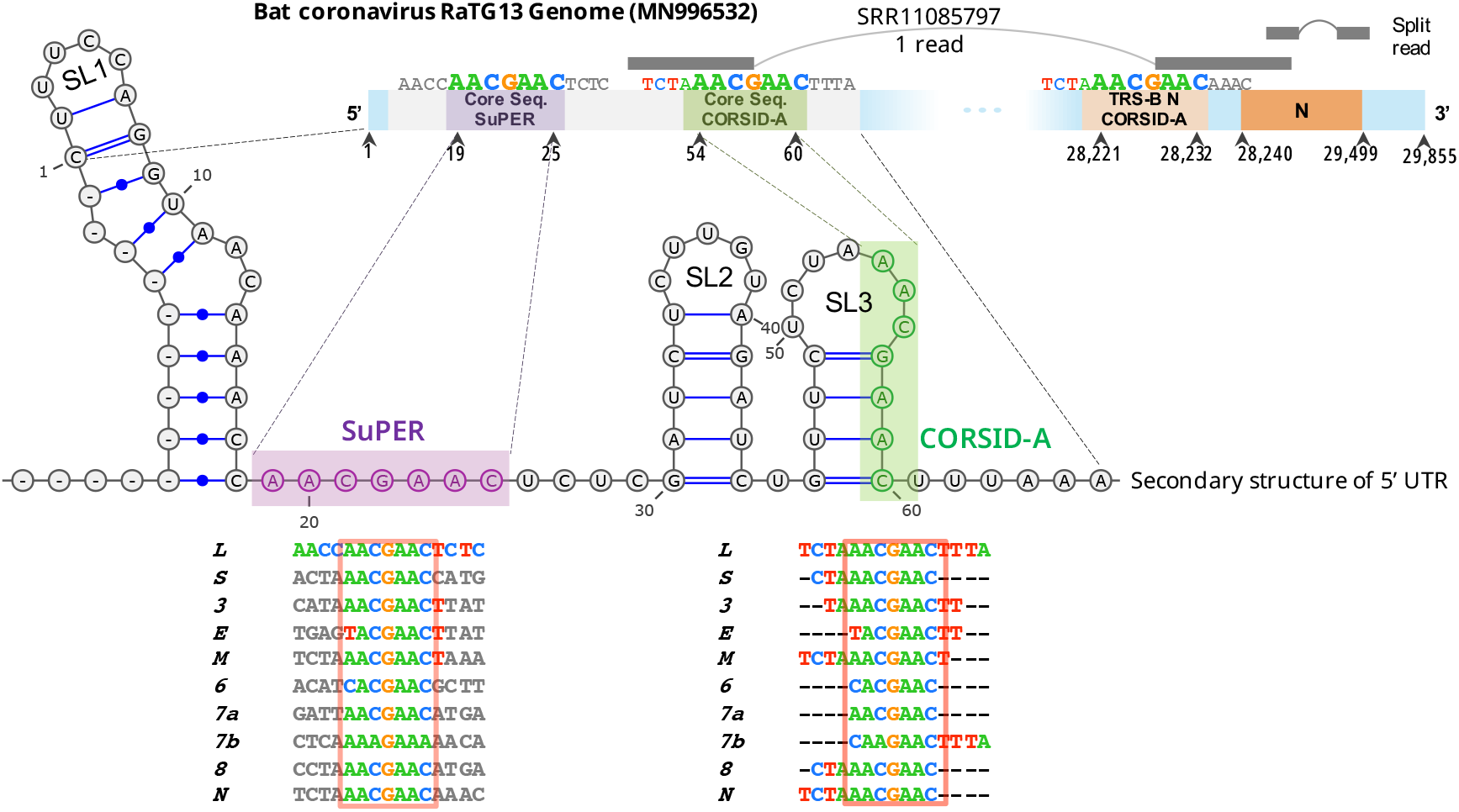
CORSID-A finds the correct TRS-L site in a *Sarbecovirus* unlike SuPER. Although SuPER uses a hard-coded motif to identify core sequences, it incorrectly identified TRS-L (positions 19 −25) in genome MN996532 (Bat coronavirus RaTG13). By contrast, CORSID-A found the TRS-L at a different location (54 − 60). We verified CORSID-A’s position using the corresponding RNA-seq data (SRR11085797). We aligned the reads to the reference genome using a splice-aware aligner, STAR [6]. The resulting alignment had a single split read, spanning positions 54 and 28221, which matches CORSID-A’s TRS-L and the TRS-B for gene *N*. Moreover, TRS-L sites in sarbecoviruses occur in stem loop (SL3) [4], which co-incides with the TRS-L site identified by CORSID-A.

**Figure S3:**
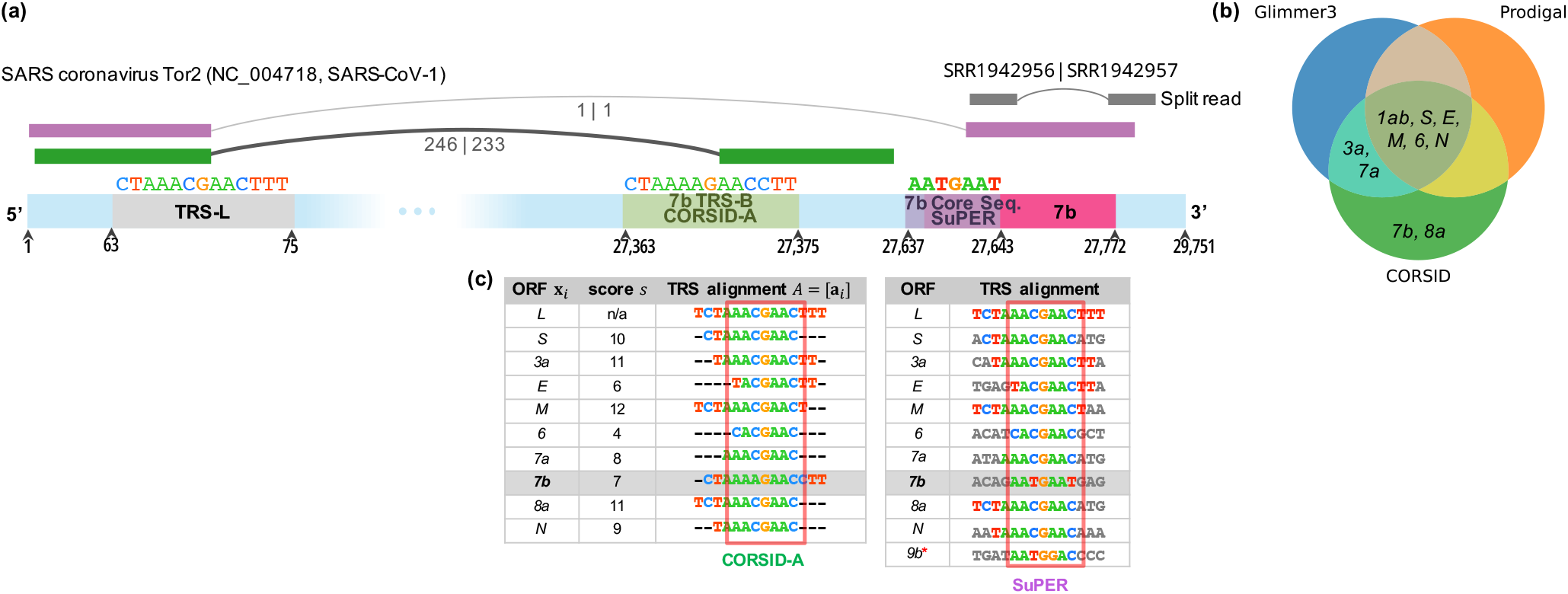
RNA-seq data supports the TRS-B found by CORSID-A, and CORSID identifies more genes than Glimmer3 and Prodigal in a SARS-CoV-1 genome (NC 004718) (a) For SARS-CoV-1 (NC 004718), CORSID-A identified a TRS-B corresponding to *ORF7b* supported by 246 and 233 split reads in RNA-sequencing samples SRR1942956 and SRR1942957, respectively. For the same genome, SuPER identified a TRS-B site that is supported by only one read in each RNA-sequencing sample. (b) A Venn diagram of the genes correctly identified by the three methods. (c) The TRS alignment of CORSID-A, and aligned core sequence with flanking regions of SuPER. “*”: SuPER identified a TRS-B for *9b* but its start codon is located at the second to fourth nucleotide of the core sequence, and the Hamming distance is 2. CORSID-A did not find this TRS-B as it occurs outside the candidate region of *9b*. Moreover, in previous studies *ORF9b* has been hypothesized to be translated via a ribosomal leaky scanning mechanism [19, 23, 27], explaining the absence of an associated TRS-B site.

**Figure S4:**
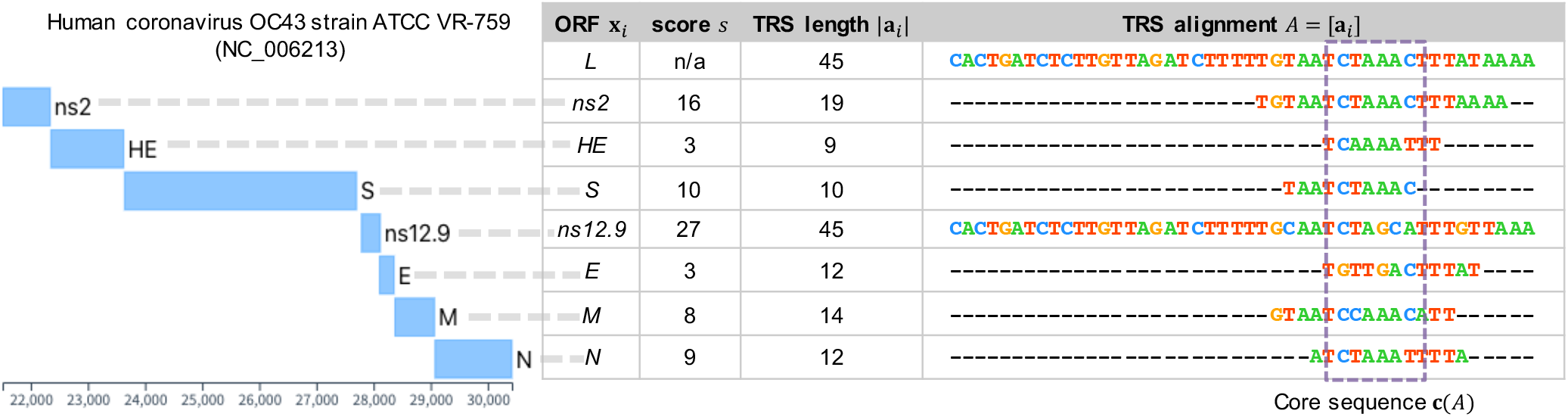
Location of genes and the TRS alignment inferred by CORSID for Human coronavirus OC43 strain ATCC VR-759 (genome NC 006213). This genome has the longest TRS-B with a length of 45 nucleotides (with only 6 mismatches) upstream of gene *ns12.9*, indicative of a recombination event.

**Figure S5:**
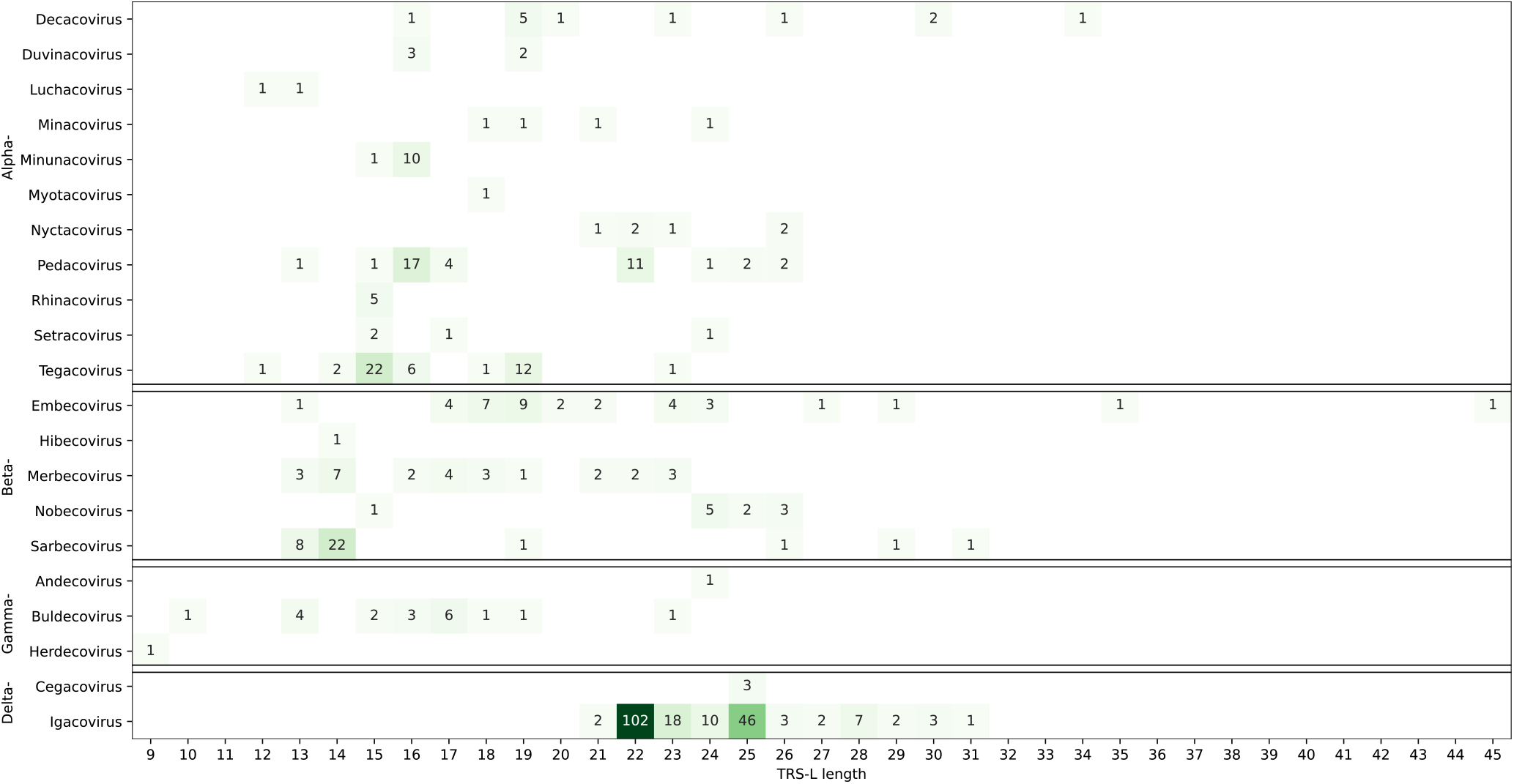
Several coronaviruses of each genus have TRS-L regions much longer (median: 22 nt) than the core sequence length (6 − 7 nt) indicating recombination events.

**Figure S6:**
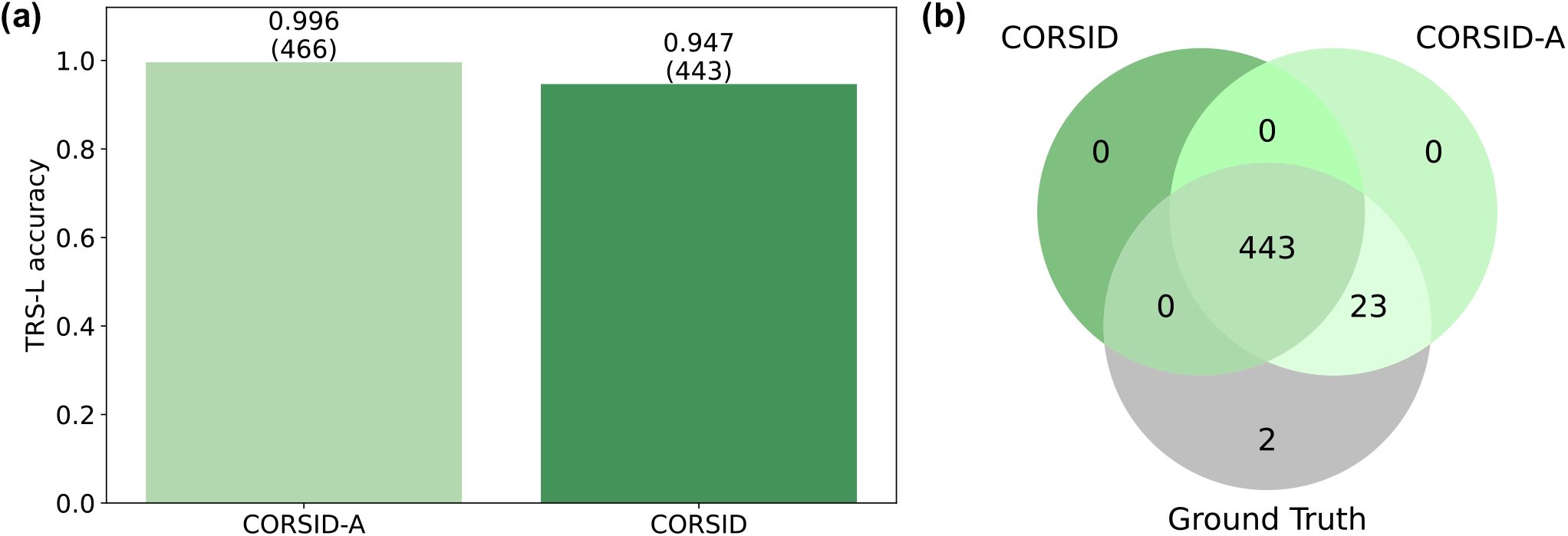
CORSID shows modest reduction in TRS-L accuracy compared to CORSID-A. (a) TRS-L accuracy of CORSID and CORSID-A. (b) Venn diagram of the TRS-L identified by CORSID and CORSID-A compared to the ground truth. Both methods failed on genomes MK211372 and MK472070 (discussed in Appendix B.4).

**Figure S7:**
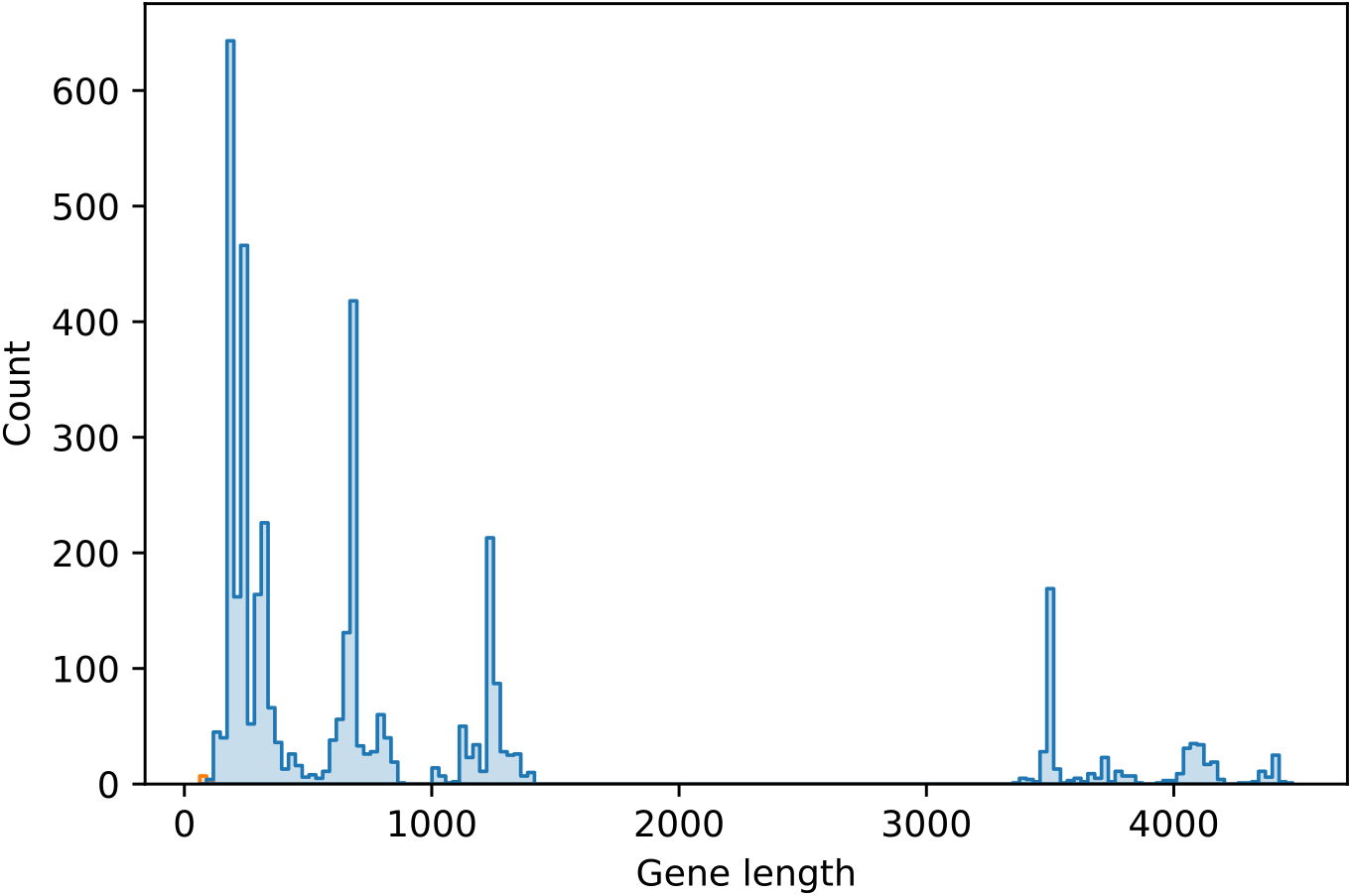
Coronavirus genes are almost always longer than 100 nt. Histogram of annotated gene lengths from 468 genomes (*ORF1ab* not included). Only 10 out of the 3637 genes are shorter than 100 nt.

**Figure S8:**
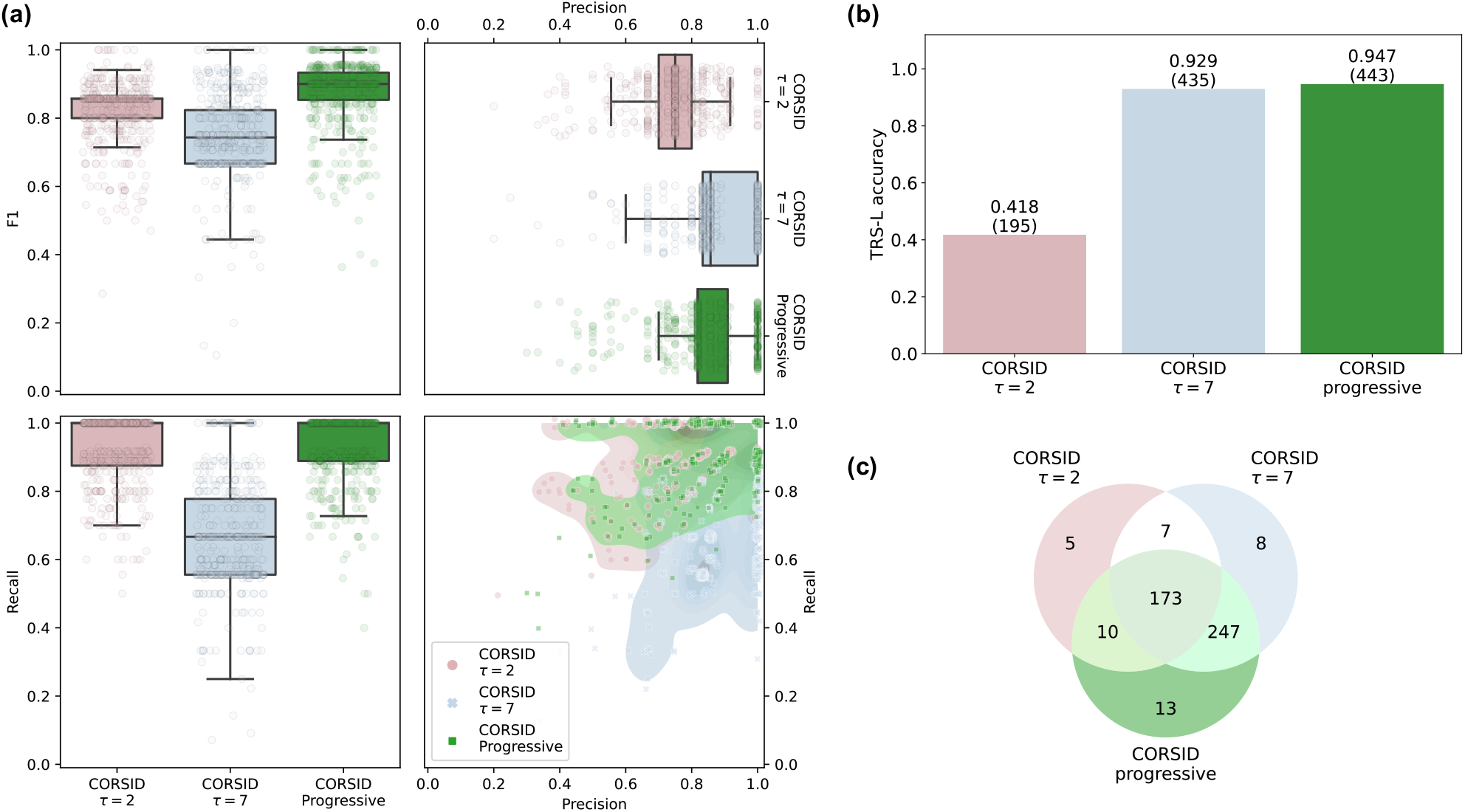
Using the progressive approach rather than directly solving *τ* = *τ*_max_ = 7 or *τ* = *τ*_min_ = 2 leads to better performance. (a) The *F*_1_ score (harmonic mean between precision and recall) is shown in top left panel. We show the precision and recall in the top right and lower left panel, respectively, and the joint distribution in the lower right panel. (b) TRS-L accuracy of setting the minimum matching score threshold to *τ* = 2 and *τ* = 7, compared with the progressive approach. (c) Venn diagram of genome sets with correctly identified TRS-L by the three versions of CORSID.

**Figure S9:**
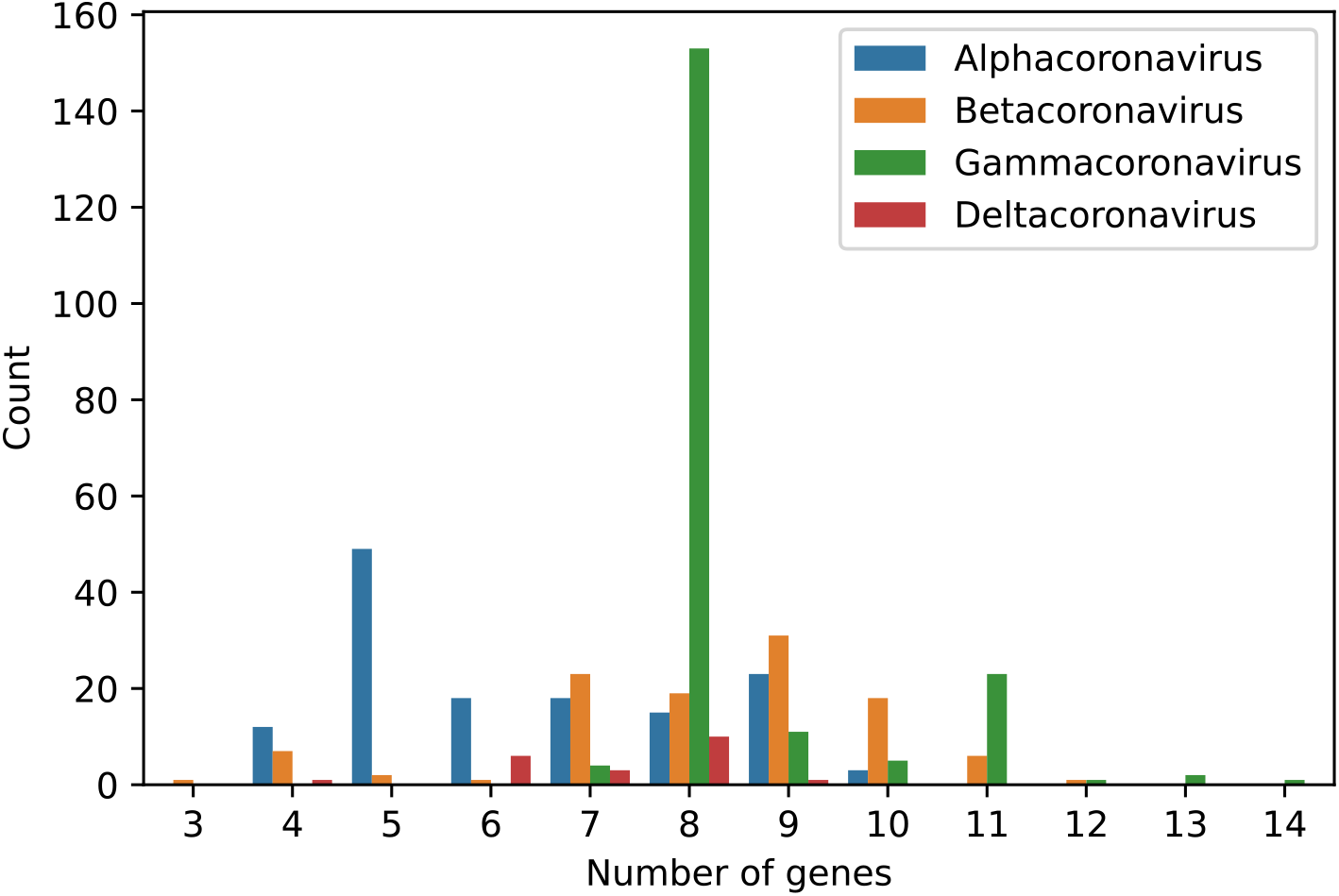
Number of annotated genes varies across the four genera of coronaviruses. (Median: 8, min: 3, max: 14).

**Figure S10:**
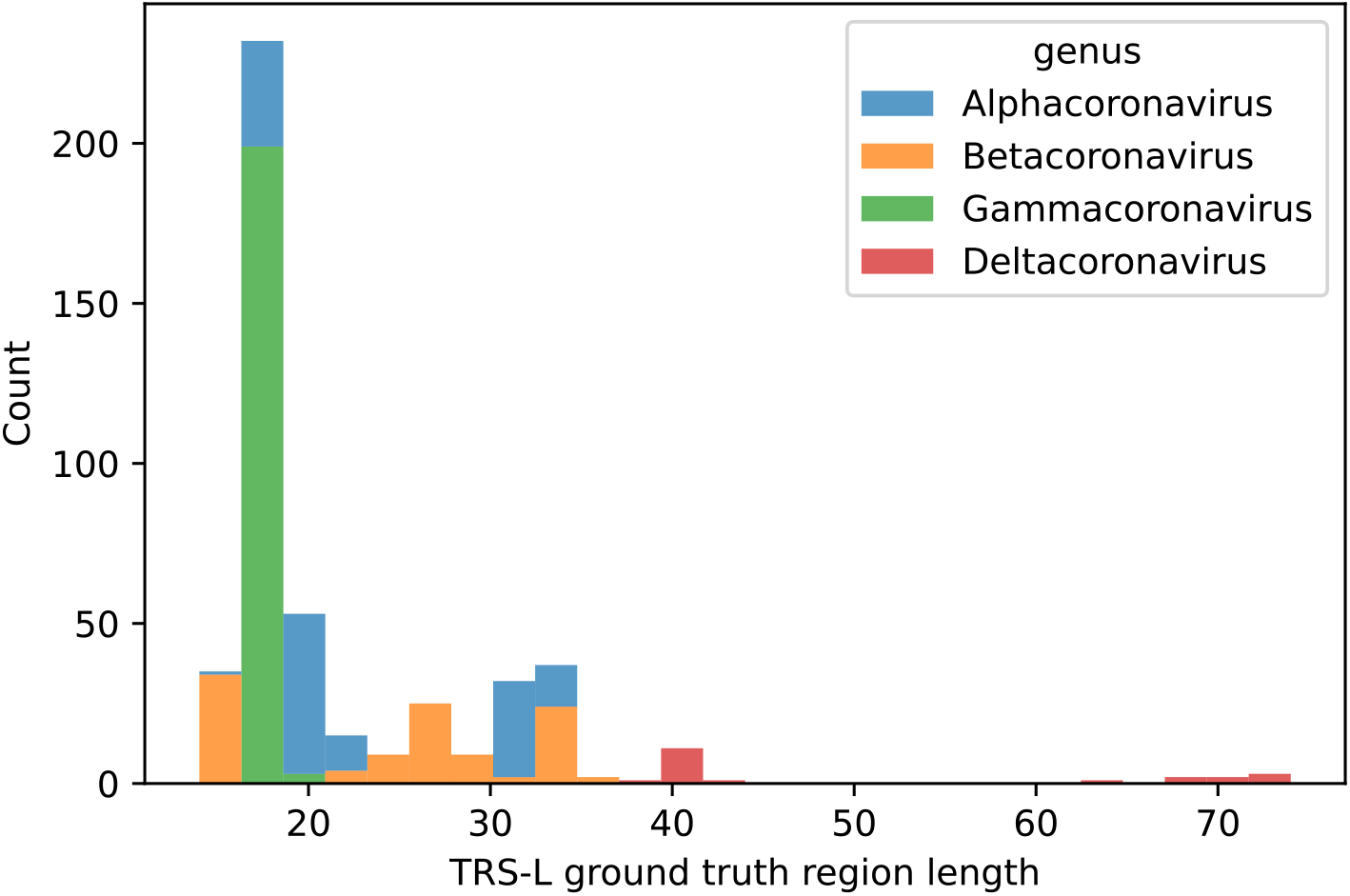
Length of the TRS-L region varies across the four genera of coronaviruses.

**Figure S11:**
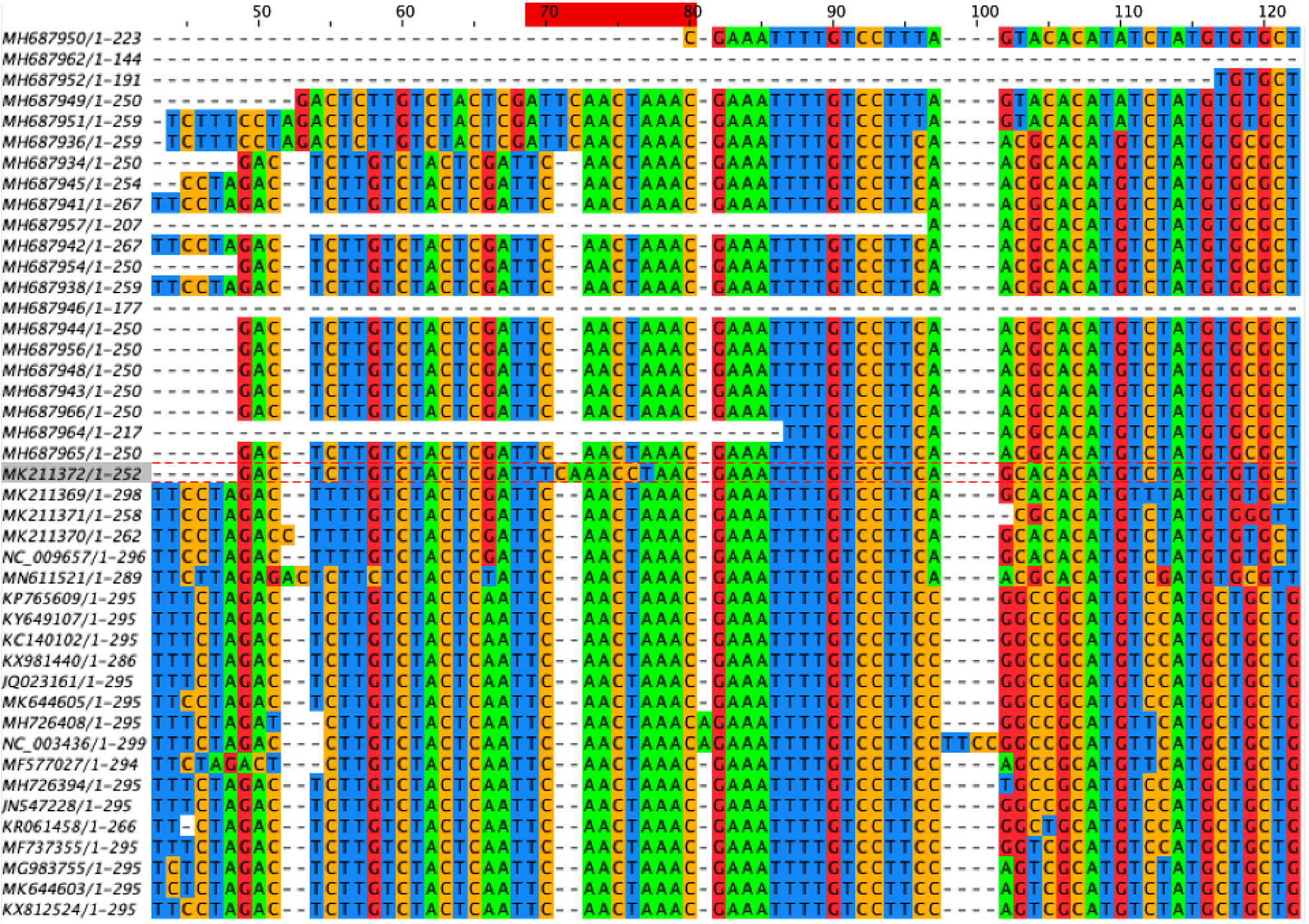
MSA of leader regions of pedacoviruses. We highlight the MK211372 genome and indicate the TRS-L region with a red bar on top. MK211372 differs from other sequences since it contains multiple indels in the TRS-L region. As such, the TRS-L region cannot be found in this genome.

**Figure S12:**
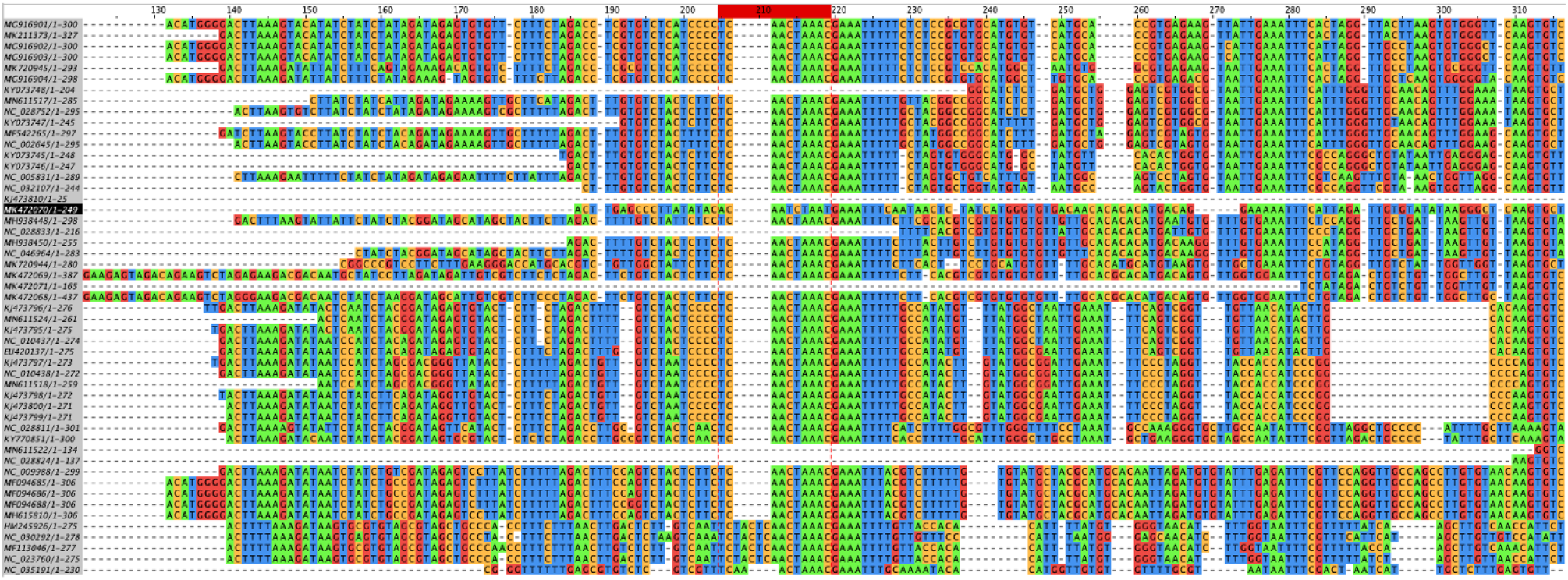
MSA of leader regions of some alphacoronaviruses. We highlight MK472070 genome and indicate the TRS-L region with red bar on top. MK472070 differs from others since it contains multiple indels in the TRS-L region. As such, the TRS-L region cannot be found in this genome.

**Table S2:** Number of coronaviruses of each genus and subgenus included and exlcuded in this study.

| **Genus** | **Subgenus** | **# Included genomes** | **# Excluded genomes** |
| --- | --- | --- | --- |
| Alphacoronavirus | Colacovirus | 0 | 2 |
|  | Decacovirus | 12 | 2 |
|  | Duvinacovirus | 5 | 1 |
|  | Luchacovirus | 2 | 0 |
|  | Minacovirus | 4 | 2 |
|  | Minunacovirus | 11 | 0 |
|  | Myotacovirus | 1 | 0 |
|  | Nyctacovirus | 6 | 2 |
|  | Pedacovirus | 39 | 6 |
|  | Rhinacovirus | 5 | 2 |
|  | Setracovirus | 4 | 0 |
|  | Tegacovirus | 45 | 1 |
|  | Unclassified | 4 | 1 |
| Subtotal |  | 138 | 19 |
| Betacoronavirus | Embecovirus | 36 | 1 |
|  | Hibecovirus | 1 | 0 |
|  | Merbecovirus | 27 | 1 |
|  | Nobecovirus | 11 | 0 |
|  | Sarbecovirus | 34 | 6 |
| Subtotal |  | 109 | 8 |
| Deltacoronavirus | Andecovirus | 1 | 0 |
|  | Buldecovirus | 19 | 0 |
|  | Herdecovirus | 1 | 0 |
| Subtotal |  | 21 | 0 |
| Gammacoronavirus | Cegacovirus | 3 | 0 |
|  | Igacovirus | 196 | 10 |
|  | Unclassified | 1 | 0 |
| Subtotal |  | 200 | 10 |
| Totals |  | 468 | 37 |

